# Overlapping roles of Arabidopsis INCURVATA11 and CUPULIFORMIS2 as Polycomb Repressive Complex 2 accessory proteins

**DOI:** 10.1101/2024.03.15.585069

**Authors:** Riad Nadi, Lucía Juan-Vicente, Samuel Daniel Lup, Yolanda Fernández, Vicente Rubio, José Luis Micol

**Author notes:** These authors contributed equally to this work (Riad Nadi, Lucía Juan-Vicente). Corresponding author: J.L. Micol.

## Abstract

Polycomb Repressive Complex 2 (PRC2) catalyzes the trimethylation of lysine 27 of histone H3 (H3K27me3) and plays a key role in epigenetic repression of gene expression in plants and animals. PRC2 core components have all been identified in *Arabidopsis thaliana*, with an expanding list of accessory proteins, some of which facilitate the recruitment of PRC2 to specific targets. INCURVATA11 (ICU11) is a 2-oxoglutarate and Fe^2+^-dependent dioxygenase that was previously shown to be a likely PRC2 accessory protein. In Tandem Affinity Purification (TAP)-based screens for interacting partners of ICU11 and its redundant paralog CUPULIFORMIS2 (CP2), we discovered that ICU11 interacts with four PRC2 core components, including EMBRYONIC FLOWER 2 (EMF2), and with the accessory proteins EMF1, TELOMERE REPEAT BINDING 1 (TRB1), TRB2, and TRB3. CP2 did not interact with PRC2 core components, nor with TRB1, TRB2, or TRB3, but did interact with TRB4 and TRB5. Both ICU11 and CP2 interacted with the nuclear proteins NAC DOMAIN CONTAINING PROTEIN 50 (NAC050), NAC052 and COP9 SIGNALOSOME SUBUNIT 1 (CSN1). Bimolecular Fluorescence Complementation (BiFC) assays revealed that ICU11 and CP2 both interact with the PRC2 core components CURLY LEAF and SWINGER, and the accessory proteins LIKE HETEROCHROMATIN PROTEIN 1, TRB1, and TRB3. ICU11 and CP2 did not interact with each other. Beyond their phenotypes, transcriptomic profiles revealed strong similarities between *emf2-3* and the double mutant *icu11-5 cp2-1*, as well as with mutants in PRC2 core components. A significant proportion of the genes mis-regulated in *icu11-5 cp2-1* are known to harbor H3K27me3 repressive marks in the wild type. Our results provide further evidence that ICU11 acts as a PRC2 accessory protein, and strongly suggest that CP2 plays a similar role.

## INTRODUCTION

The first gene encoding a Polycomb group (PcG) protein was discovered by the characterization of a mutation in the fruit fly *Drosophila melanogaster* (Lewis, 1947). PcG proteins are highly conserved among eukaryotes and epigenetically repress the expression of genes controlling growth, development, and environmental adaptation (Jiao et al., 2020; Xiao and Wagner, 2015). PcG proteins form part of two heteromultimeric Polycomb Repressive Complexes (PRCs), which perform different epigenetic activities: PRC1 is a histone H2A ubiquitin ligase, whereas PRC2 is a histone H3 lysine 27 (H3K27) methyltransferase (Bratzel et al., 2010).

Plant PRCs function in many critical developmental stages and events, such as the transition from embryo to seedling (Bouyer et al., 2011), gametophyte and seed development (Roszak and Köhler, 2011), and vernalization and flowering induction (Pazhouhandeh et al., 2011; Tian et al., 2019; Whittaker and Dean, 2017). In Arabidopsis (*Arabidopsis thaliana*), PRC2 comprises eight core components: CURLY LEAF (CLF: Goodrich et al., 1997), SWINGER (SWN; Chanvivattana et al., 2004), MEDEA (MEA; Grossniklaus et al., 1998), FERTILIZATION INDEPENDENT SEED 2 (FIS2; Luo et al., 1999), EMBRYONIC FLOWER 2 (EMF2; Yoshida et al., 2001), VERNALIZATION2 (VRN2; Gendall et al., 2001), FERTILIZATION-INDEPENDENT ENDOSPERM (FIE; Ohad et al., 1999), and MULTICOPY SUPRESSOR OF IRA (inhibitory regulator of the RAS-cAMP pathway) 1 (MSI1; Hennig et al., 2003; Köhler et al., 2003). The Arabidopsis PRC1 core components include three B Lymphoma Mo-MLV Insertion Region 1 (BMI1) homologs (BMI1A, BMI1B, and BMI1C) and two RING FINGER proteins (RING1A and RING1B) (Sanchez-Pulido et al., 2008).

Accessory proteins facilitate the recruitment of PRC1 and PRC2 to specific chromatin regions; for example, LIKE HETEROCHROMATIN PROTEIN 1 (LHP1) and the plant-specific protein EMBRYONIC FLOWER 1 (EMF1) interact with each other and with PRC1 and PRC2 core components (Bratzel et al., 2010; Calonje et al., 2008; Derkacheva et al., 2013; Mozgova and Hennig, 2015). LHP1 contributes to the maintenance of the tri-methylated H3K27 (H3K27me3) chromatin repressive state through the continuous recruitment of PRC2 to regions enriched with the H3K27me3 mark (Ramirez-Prado et al., 2019; Turck et al., 2007; Zhang et al., 2007). EMF1 contributes to H3K27me3 deposition at a subgroup of PRC2 target genes, and is also required for histone H2A monoubiquitination by PRC1 (Kim et al., 2012).

Lack of vegetative development and the formation of flower-like organs immediately after germination—the so-called embryonic flowers—is a conspicuous phenotype that was first observed in *emf1* mutants (Aubert et al., 2001; Sung et al., 1992; Yang et al., 1995; Yoshida et al., 2001). Embryonic flowers are also produced by the *telomere repeat binding1-2* (*trb1-2*) *trb2-1 trb3-2* triple mutant (Yang et al., 2013; Zhou et al., 2018). Arabidopsis TRB1, TRB2, and TRB3 bind to telomeric repeat DNA sequences to maintain chromosome ends (Klepikova et al., 2016; Lee and Cho, 2016; Nadi et al., 2023; Schubert et al., 2006), and are thought to recruit the PRC2 complex to certain genes for H3K27me3 deposition (Zhou et al., 2018).

The 2-oxoglutarate and Fe(II)-dependent dioxygenase (2OGD, also called 2ODD) domain characterizes the second largest protein superfamily in the plant kingdom (Martinez and Hausinger, 2015) and is represented by about 150 genes in Arabidopsis (Kawai et al., 2014; Nadi et al., 2018). 2OGD proteins catalyze oxidative reactions using 2-oxoglutarate (also called α-ketoglutarate) and molecular oxygen as co-substrates, and Fe^2+^ as a cofactor (Islam et al., 2018). Phylogenetic analyses of plant 2OGDs grouped them into the DOXA, DOXB, DOXC, and JUMONJI (JMJ) protein classes, with demonstrated functions that include DNA and RNA demethylation, collagen hydroxylation, a diverse range of metabolic processes, and histone demethylation, respectively (Islam et al., 2018; Kawai et al., 2014). Two Arabidopsis DOXB-type 2OGDs, INCURVATA11 (ICU11) and CUPULIFORMIS2 (CP2), are redundant components of the epigenetic machinery (Mateo-Bonmatí et al., 2018; Nadi et al., 2023). Whereas *icu11* mutants exhibit a mild morphological phenotype consisting of hyponastic leaves and early flowering, and *cp2* mutants are phenotypically wild type, *icu11 cp2* double mutants skip vegetative development and develop embryonic flowers immediately after germination, culminating in plant death 20–40 days after stratification (Mateo-Bonmatí et al., 2018; Nadi et al., 2023).

Based on co-immunoprecipitation (co-IP) analyses, ICU11 was proposed to be a PRC2 accessory protein, probably involved in H3K36me3 demethylation at the *FLOWERING LOCUS C* (*FLC*) floral repressor gene (Bloomer et al., 2020). Here, we provide further evidence for ICU11 as a PRC2 accessory protein through experimental approaches complementary to co-IP, including an *in vitro* tandem affinity purification (TAP)-based screen and *in vivo* heterologous bimolecular fluorescence complementation (BiFC) assays. We also used these approaches to identify several interacting partners of CP2, some of which are PRC2 core components or accessory proteins. Furthermore, using RNA sequencing (RNA-seq), we identified many genes that are upregulated in the lethal embryonic flowers of the *icu11-5 cp2-1* double mutant and involved in flower development, as previously shown by microarray analysis in the *emf2-3* single mutant (Kim et al., 2010). Taken together, our results confirm that ICU11 is a PRC2 accessory protein and strongly suggest that CP2 also plays this role.

## RESULTS

### ICU11 interacts with PRC2 core components and accessory proteins in a TAP-based screen

To identify interactors of ICU11 and CP2, we carried out a TAP-based screen followed by liquid chromatography electrospray ionization and tandem mass spectrometry (LC-ESI-MS/MS). Specifically, we used C-terminal translational fusions of ICU11 and CP2 to the GS^Rhino^ tag, consisting of protein G, a streptavidin-binding peptide, and rhinovirus 3C protease cleavage sites. We transformed PSB-D Arabidopsis cell cultures with *Agrobacterium tumefaciens* carrying the aforehead mentioned translational fusions. In line with previous results obtained using co-IP assays followed by tandem mass spectrometry (Bloomer et al., 2020), we determined that ICU11 interacts with the PRC2 core components EMF2, FIE, SWN, and MSI1, as well as the PRC2 accessory proteins EMF1, TRB1, TRB2, and TRB3 (Supplemental Figure S1, Supplemental Table S1 and Supplemental Dataset DS1). TRB1, TRB2, and TRB3 are components of the PWWPs-EPCRs-ARIDs-TRBs (PEAT) complexes that recruit PRC2 to telobox-related motifs present at telomeres (Tan et al., 2018; Zhou et al., 2016; Zhou et al., 2018). We also identified MSI1 and SWN as interactors of ICU11 in one of our TAP-based replicates (Supplemental Dataset DS1). In contrast to Bloomer et al. (2020), we did not identify CLF or LHP1 as ICU11 interactors.

Among the best-represented interactors of ICU11, we also noticed three paralogous proteins, which are predicted to be nuclear: the AT5G66000 hypothetical protein and the AT3G17460 and AT4G35510 uncharacterized proteins with a PHD finger domain. AT5G66000 was previously detected as an interactor of EMF1, CLF, and ICU11 in the co-IP assays performed by Bloomer et al. (2020), in which these authors also detected AT3G17460 as an ICU11 interactor, and AT4G35510 as an interactor of CLF but not ICU11.

### CP2 interacts with TRB4, TRB5, and other nuclear proteins in a TAP-based screen

We also performed a TAP-based screen to identify CP2 interactors, from which we detected no PRC2 core component. CP2 strongly interacted with TRB4 and TRB5 (Supplemental Figure S1 and Supplemental Table S1), two poorly characterized members of the TRB family; however, their TRB1, TRB2, and TRB3 paralogs were not detected as CP2 interactors. We excluded any possible ambiguity in the interactions of ICU11 and CP2 with the TRB proteins by checking that the peptides identified from each TRB were protein-specific, the only exception being one peptide whose sequence matches an identical region in TRB2 and TRB3 (Supplemental Figure S3). Other nuclear proteins identified as interactors of CP2 but not of ICU11 were DEVELOPMENT RELATED MYB-LIKE1 (DRMY1) and DRMY PARALOG 1 (DP1; Supplemental Figure S1 and Supplemental Table S1), which belong to the single repeat MYB family of transcription factors. Whereas the *dp1* mutant is indistinguishable from the wild type, *drmy1* loss-of-function mutants exhibit pleotropic defects in root, vegetative, and floral development, but not in flowering time (Wu et al., 2018; Yanhui et al., 2006; Zhu et al., 2020). Another interactor of CP2 but not of ICU11 was INOSITOL REQUIRING 80 (INO80; Supplemental Figure S1 and Supplemental Table S1), a nuclear chromatin remodeling factor conserved among eukaryotes. Depletion of INO80 represses photomorphogenesis and causes multiple developmental defects including reduced plant size, late flowering, abnormal shape of reproductive organs, reduced pollen grain number per anther, and smaller siliques (Kang et al., 2019; Yang et al., 2020; Zhang et al., 2015).

Another CP2 interactor we identified was the JMJ-type 2OGD protein INCREASE IN BONSAI METHYLATION 1 (IBM1; Supplemental Figure S1 and Supplemental Table S1), a known H3K9me2/1 demethylase (Miura et al., 2009). Several *ibm1* alleles perturb leaf and flower morphogenesis and reduce fertility. The depletion of IBM1 increases H3K9me marks and DNA methylation in the CHG and CHH genomic contexts (Saze et al., 2008).

### Nuclear proteins that interact with both ICU11 and CP2 in TAP-based screens

Our TAP assays also revealed nuclear proteins that interact with both ICU11 and CP2 (Supplemental Figure S1, Supplemental Table S1 and Supplemental Dataset DS1). Two of these were the paralogous transcription factors NAC050 and NAC052, which associate with the histone demethylase JMJ14 in the negative regulation of flowering through the removal of the H3K4me3 mark at flowering regulator genes such as *FLC* (Ning et al., 2015). Neither NAC050 nor NAC052 was identified as an ICU11 interactor by Bloomer et al. (2020). By contrast, JMJ14 was identified as an ICU11 interactor by Bloomer et al. (2020) but was not detected in our TAP assays.

We also identified COP9 SIGNALOSOME SUBUNIT 1 (CSN1), also named FUSCA 6 (FUS6), a member of the CONSTITUTIVE PHOTOMORPHOGENESIS 9 (COP9) signalosome complex, which maintains skotomorphogenesis by repressing photomorphogenesis (Qin et al., 2020; Wang et al., 2002), and is required for the proper development of floral organs (Wang et al., 2003). CSN1 was not detected by Bloomer et al. (2020). Neither of our two TAP-based screens revealed any interaction between ICU11 and CP2, despite their shared interactors.

### ICU11 and CP2 interact with PRC2 core components and accessory proteins in BiFC assays

To obtain complementary evidence for the results of our TAP-based screens, we performed heterologous BiFC assays through the transient transformation of *Nicotiana benthamiana* leaves (Kerppola, 2006; Martin et al., 2009). The co-infiltration of such leaves with constructs encoding the N-terminal half of enhanced yellow fluorescence protein (EYFP) fused to ICU11 (nEYFP-ICU11) and the C-terminal half of EYFP fused to CLF (cEYFP-CLF), SWN (cEYFP-SWN), or LHP1 (cEYFP-LHP1) all produced strong nuclear EYFP signals (Figure 1A–I), which is consistent with the known nucleoplasmic colocalization of ICU11 (Mateo-Bonmatí et al., 2018), CLF (Schubert et al., 2006), SWN (Wang et al., 2006), and LHP1 (Zemach et al., 2006). We established that nEYFP-CP2 interacts with cEYFP-CLF, cEYFP-SWN, and cEYFP-LHP1 (Figure 1P–X). All co-infiltrations of nEYFP-ICU11 or nEYFP-CP2 with cEYFP-TRB1 or cEYFP-TRB3 resulted in strong EYFP signals (Figure 1J–O, Y–AD), consistent with the known subnuclear localization of TRB1 and TRB3 (Zhou et al., 2016). As a positive control, both coinfiltrations of cEYFP-LHP1 with nEYFP-UBP12 and nEYFP-UBP13 rendered strong nuclear signals, as previously described (Supplemental Figure S3A-F; Derkacheva et al., 2016). The absence of interaction between ICU11 and CP2 was confirmed by coinfiltration of *Nicotiana benthamiana* leaves with nEYFP-ICU11 and cEYFP-CP2 (Supplemental Figure S3G-I). nEYFP-ICU11 and nEYFP-CP2 were used as negative controls (Supplemental Figure S3O-Q).

**Figure 1.**
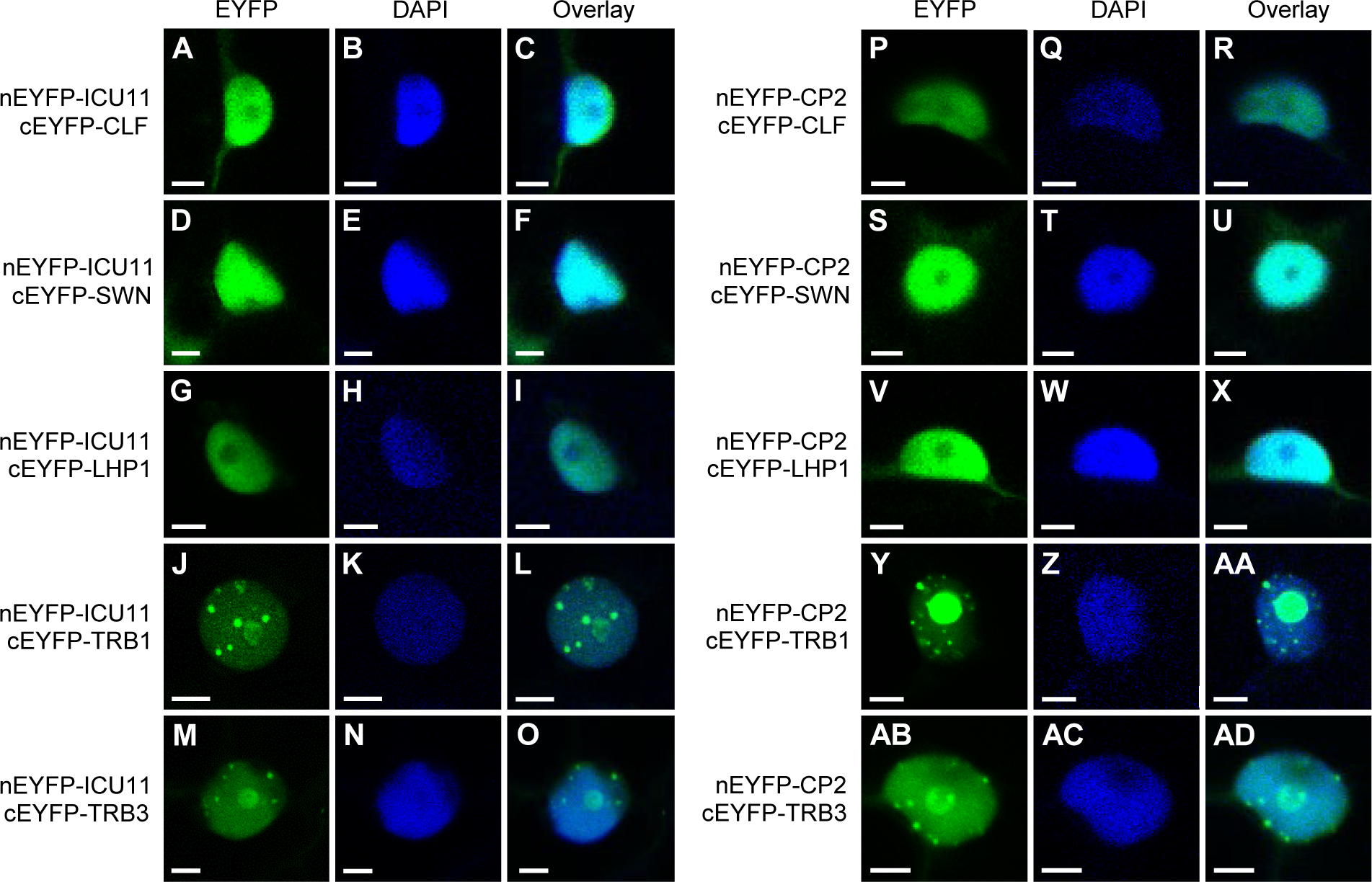
*In vivo* interactions of ICU11 and CP2 with proteins with known epigenetic roles. Bimolecular fluorescence complementation assays showing interaction between the indicated proteins. Individual nuclei of *Nicotiana benthamiana* leaves co-infiltrated with the constructs *nEYFP-ICU11* or *nEYFP-CP2* with *cEYFP-CLF*, *cEYFP-LHP1*, *cEYFP-TRB1*, or *cEYFP-TRB3*. Fluorescent signals correspond to EYFP (A, D, G, J, M, P, S, V, Y, AB), DAPI (B, E, H, K, N, Q, T, W, Z, AC), and their overlay (C, F, I, L, O, R, U, X, AA, AD). Scale bars, 5 µm.

### The transcriptomic profile of the *icu11-5 cp2-1* double mutant resemble that of the *emf2-3* single mutant

We previously used both RNA-seq and reverse transcription-quantitative PCR (RT-qPCR) to show that the *icu11-1* mutant, in the Arabidopsis S96 genetic background, transcriptionally misregulates hundreds of genes (Mateo-Bonmatí et al., 2018). As *ICU11* and *CP2* encode putative PRC2 accessory proteins and the morphological phenotype of their double mutant combinations (namely, embryonic flowers) resembles that of the *emf1* and *emf2* single mutants, we compared the transcriptome of the *icu11-5 cp2-1* double mutant with that of the PRC2 strong loss-of-function mutant *emf2-3*. We used clustered regularly interspaced short palindromic repeat (CRISPR)/CRISPR-associated nuclease 9 (Cas9)-mediated mutagenesis to obtain the *icu11-5* and *icu11-6* alleles of *ICU11* in a Col-0 genetic background to avoid possible differences in gene expression due to the genetic background (Nadi et al., 2023).

We performed RNA-seq analyses of the Col-0, *icu11-5* and *cp2-1* seedlings, and the *icu11-5 cp2-1* and *emf2-3* embryonic flowers, which were all collected 10 days after stratification (das); Col-0 inflorescences were also collected 40 das for RNA-seq (Figure 2, Supplemental Table S2 and Supplemental Dataset DS2). We only identified 23 upregulated genes and 5 downregulated genes in the *cp2-1* mutant relative to Col-0; the morphological phenotype of this mutant is indistinguishable from the wild type. We also identified 738 upregulated genes and 78 downregulated genes in the *icu11-5* mutant, whose morphological phenotype is relatively weak. By contrast, the number of de-regulated genes in the *icu11-5 cp2-1* and *emf2-3* embryonic flowers was similar: these plants showed 3199 upregulated genes and 1770 downregulated genes, and 2520 upregulated genes and 1774 downregulated genes, respectively, when compared to Col-0 seedlings. Moreover, the Col-0 inflorescences showed the expected strong transcriptomic differences when compared to Col-0 seedlings, with 5431 upregulated genes and 3084 downregulated genes, in agreement with similar data previously published (Klepikova et al., 2016).

**Figure 2.**
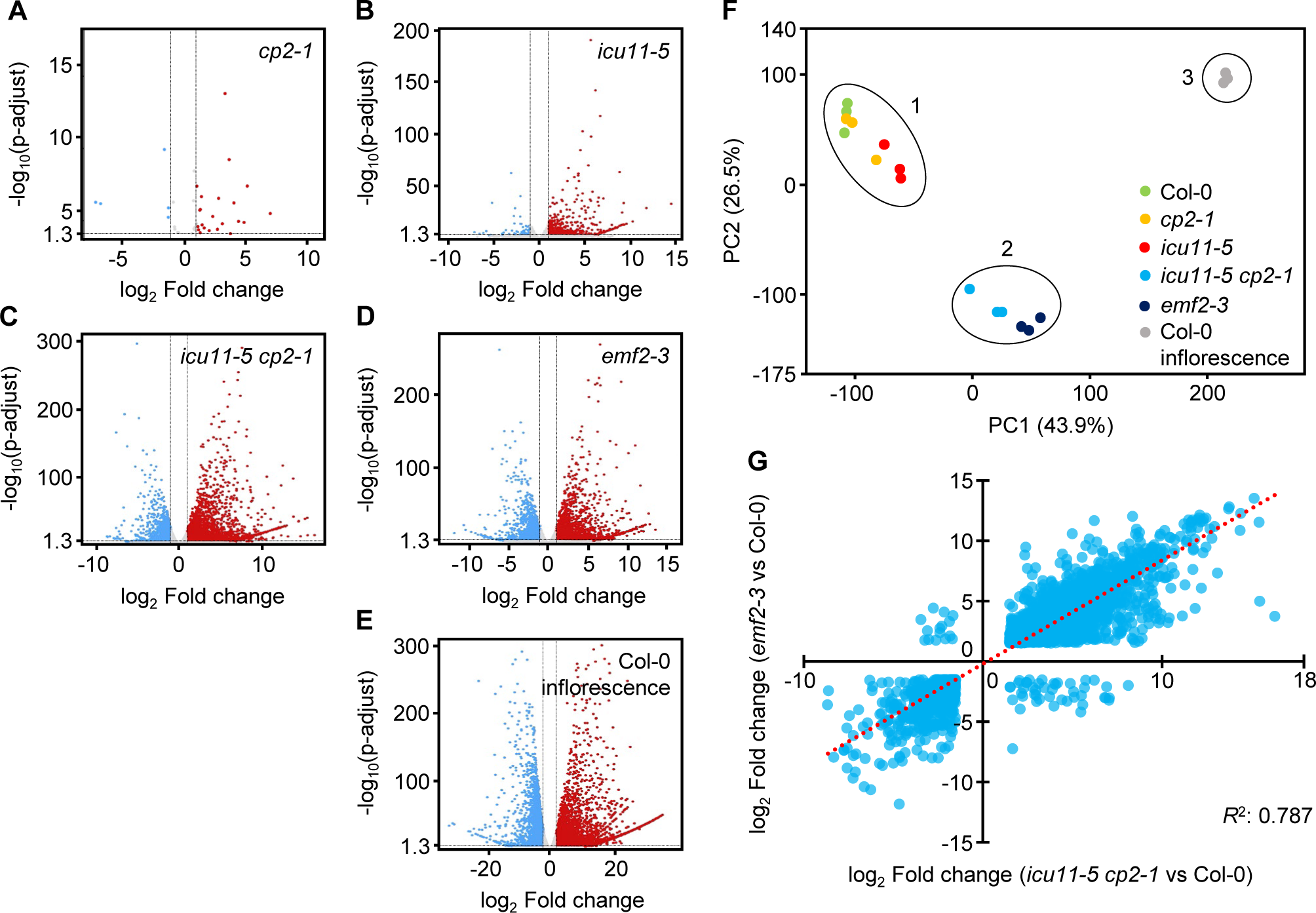
Transcriptomic profiling of *icu11-5 cp2-1* and *emf2-3* embryonic flowers. (A–E) Volcano plots representing differentially expressed genes (DEGs) in *cp2-1* (A) and *icu11-5* (B) seedlings; *icu11-5 cp2-1* (C) and *emf2-3* (D) embryonic flowers; and Col-0 (E) inflorescences, all compared to Col-0 seedlings. Blue and red dots indicate significantly downregulated and upregulated genes, respectively, with a Benjamini and Hochberg corrected *p-*value < 0.05. Total RNA was extracted from three biological samples collected 10 (A–D) or 40 (E) das. (F) Principal component analysis of the transcriptomic profiles showing three clusters: (1) Col-0, *cp2-1,* and *icu11-5* seedlings; (2) *icu11-5 cp2-1,* and *emf2-3* embryonic flowers; and (3) Col-0 inflorescences. Each dot represents a biological replicate. (G) Scatterplot showing the positive correlation between the relative expression levels of DEGs of the *icu11-5 cp2-1* double mutant and those of the *emf2-3* single mutant, both relative to Col-0 seedlings. Log_2_ values ranging from –1.5 to 1.5 were not plotted. The best-fit line is shown as a red dashed line, and the *R^2^* value is indicated.

Genes encoding MADS-box transcription factors, such as *AGAMOUS* (*AG*) and *SEEDSTICK* (*STK*), are flower organ identity genes repressed by PRC2 during the vegetative stage (Petrella et al., 2020; Schubert et al., 2006). In the present study, *AG*, *SHATTERPROOF* (*SHP2*), and *STK* were found upregulated in in *icu11-5* seedlings, *icu11-5 cp2-1* and *emf2-*embryonic flowers, and Col-0 inflorescences, but not in *cp2-1* seedlings, which we confirmed using RT-qPCR. Among the genes upregulated in *cp2-1* seedlings, we found *EARLY ARABIDOPSIS INDUCED 1* (*EARLI1*), which encodes a proline-rich family protein involved in lignin biosynthesis and flowering time control (Shi et al., 2011); *PIRIN 1* (*PRN1*), a cupin-fold protein involved in seed germination, development, and the response to abscisic acid and light (Orozco-Nunnelly et al., 2014); and *RIBONUCLEASE 1* (*RNS1*), a protein that functions in cell death and the generation of tRNA-derived fragments, which are involved in the regulation of gene expression, RNA degradation, and the inhibition of protein synthesis (Goodman et al., 2022; Megel et al., 2019) (Supplemental Figure S4).

A principal component analysis identified different patterns of transcriptional misregulation, with three main clusters: one formed by the seedlings of Col-0, *cp2-1,* and to a certain extent *icu11-5*; another one consisting of the *icu11-5 cp2-1* and *emf2-3* embryonic flowers; and the last one representing the transcriptome of the Col-0 inflorescence (Figure 2F). The transcriptomes of *icu11-5 cp2-1* and *emf2-3* were similar (*R*^2^ = 0.787; Figure 2G).

A protein domain enrichment analysis revealed that *icu11-5* seedlings*, icu11-5 cp2-1* and *emf2-3* embryonic flowers and Col-0 inflorescences share an upregulation of genes in the Mitogen-Activated Protein Kinase (MAPK) cascade, an important conserved mechanism in eukaryotes that triggers the intracellular transduction response to a range of developmental and environmental signals (Jagodzik et al., 2018; Plotnikov et al., 2011; Supplemental Datasets DS3 to DS7). The *icu11-5 cp2-1* and *emf2-3* transcriptomes also had similar Gene Ontology (GO) enrichment of biological processes profiles, among the most significant of which for the upregulated genes included response to phytohormones, abiotic stresses, and transcriptional regulation (Supplemental Datasets DS5 and DS7). We also observed that most enriched GO terms in the downregulated genes in the *icu11-5 cp2-1* and *emf2-3* embryonic flowers and the Col-0 inflorescence are related to photosynthesis, chloroplast organization and biosynthesis, and sucrose biosynthesis (Supplemental Datasets DS5 and DS7).

Regarding the protein domain enrichment analysis, the genes upregulated in *icu11-5* were enriched in those encoding proteins harboring the keratin-like (K-box) and MADS-box domains (Supplemental Dataset DS3), which are associated with the regulation of flowering time (Alvarez-Buylla et al., 2000). The same categories were also enriched in the upregulated genes of *icu11-5 cp2-1* and *emf2-3* embryonic flowers, and Col-0 inflorescences, which also encompassed 13 other categories, including Non Apical Meristem (NAM), a FAD-binding domain, and WRKY domains, which are also related to the regulation of flowering (Aida et al., 1997; Liu et al., 2008; Martignago et al., 2019; Singh et al., 2014; Spedaletti et al., 2008). The genes upregulated in *icu11-5 cp2-1* and *emf2-3* embryonic flowers were significantly enriched in genes encoding transcription factors containing the APETALA2/ETHYLENE-RESPONSIVE ELEMENT BINDING FACTOR (AP2/ERF) domain (Drews et al., 1991; Feng et al., 2020; Okamuro et al., 1997; Supplemental Datasets DS5-DS7).

### The transcriptomic profile of *icu11-5 cp2-1* resembles that of mutants affected in genes encoding PRC2 core components or accessory proteins

Venn diagrams of the Differentially Expressed Genes (DEGs) of *icu11-5* and *cp2-1* seedlings and *icu11-5 cp2-1* embryonic flowers, all compared with Col-0 seedlings, showed no overlap between the genes downregulated in *icu11-5* and *cp2-1*, and only eight genes were upregulated in both *icu11-5* and *cp2-1*. We also found that 78% and 58% of the genes upregulated and downregulated in *icu11-5*, respectively, are coregulated in the *icu11-5 cp2-1* double mutant (Figure 3A, D).

**Figure 3.**
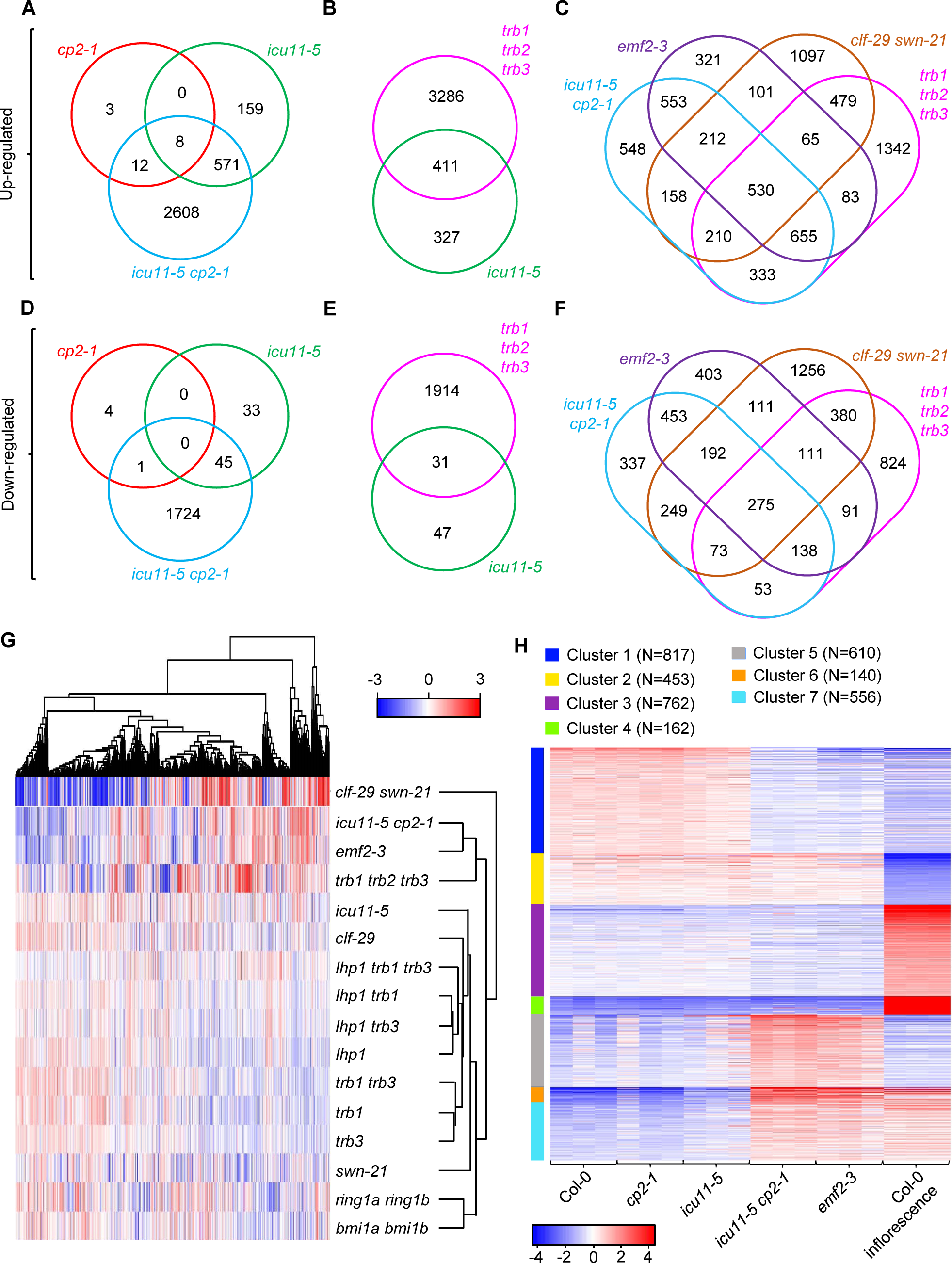
Comparison of differential expression in the *icu11-5 cp2-1* double mutant and in mutants lacking function of PRC2 core components or accessory proteins. (A–F) Venn diagrams showing the overlap between upregulated (A–C) and downregulated (D–F) genes in *cp2-1* and *icu11-5* seedlings and *icu11-5 cp2-1* embryonic flowers (A, D), *icu11-5* seedlings and *trb1 trb2 trb3* embryonic flowers (B, E), and *icu11-5 cp2-1, emf2-3* and *trb1 trb2 trb3* embryonic flowers and *clf-29 swn-21* callus-like seedlings (C, F). The abbreviations *trb1, trb2, trb3,* and *lhp1* stand for the alleles *trb1-2*, *trb2-1*, *trb3-2,* and *lhp1-4,* respectively. (G) Heatmap showing the normalized log_2_ fold-change of genes misregulated in the plants studied. Genes represented in red and blue are upregulated and downregulated, respectively. (H) k-means transcriptional clustering of the genotypes under study. Seven clusters and normalized read counts of the 3500 most variable genes were used. N is the number of genes per cluster. The color scale indicates the range of normalized log_2_ fold-change of the 3500 genes.

We conducted comparative analyses of the published transcriptomic profiles of mutants carrying mutant alleles of genes encoding PRC2 core components or accessory proteins, which exhibit morphological phenotypes ranging from wild type to callus-like, as is the case for *clf-29 swn-21* (Wang et al., 2016; Yang et al., 2013). The *trb1-2 trb2-1 trb3-2* triple mutant exhibits an embryonic flower phenotype (Zhou et al., 2018); we determined that 56% of the 411 upregulated genes and 40% of the 31 downregulated genes in *icu11-5* are similarly upregulated or downregulated in *trb1-2 trb2-1 trb3-2* (Figure 3B, E). The morphological phenotypes of *trb1-2 trb2-1 trb3-2*, *emf2-3,* and *icu11-5 cp2-1* are similar, and their transcriptomic profiles included 530 and 275 genes that are similarly upregulated or downregulated, respectively. Only 548 (17%) and 337 (19%) genes were exclusively upregulated and downregulated, respectively, in *icu11-5 cp2-1* but not in *emf2-3, clf29 swn-21, or trb1-2 trb2-1 trb3-2* (Figure 3C, F).

Hierarchical clustering of the *icu11-5* and *icu11-5 cp2-1* transcriptomic profiles and those of mutants affected in genes encoding PRC2 core components and accessory components, as well as PRC1 core components, revealed that *icu11-5 cp2-1* showed a high transcriptomic similarity to *emf2-3,* and to a lesser extent with *trb1-2 trb2-1 trb3-2.* The *icu11-5* mutant clustered with the mutans affected in PcG genes with milder morphological phenotypes, such as *clf-29* (Figure 3G).

To ascertain which set of the genes misregulated in *icu11-5 cp2-1* contributes to its embryonic flower phenotype, we performed k-clustering with the 3500 most variably expressed genes and a k value of 7 (Figure 3H). Clusters 2, 3, and 4 harbored 453, 762, and 162 genes, respectively, for which the Col-0 inflorescence presented significantly different expression levels compared to the remaining samples. Cluster 2 genes were repressed in Col-0 inflorescences, suggesting that these genes are important for vegetative development. On the contrary, 924 genes from clusters 3 and 4 were highly expressed in Col-0 inflorescences, suggesting a role in reproductive development instead. We identified 817 genes from cluster 1 (downregulated) and 556 genes from cluster 7 (upregulated) as being coregulated in the *icu11-5 cp2-1* and *emf2-3* embryonic flowers and Col-0 inflorescences in comparison to Col-0 seedlings. Moreover, cluster 5 contained 610 genes that are highly expressed in *icu11-5 cp2-1* and *emf2-3* but not in the remaining samples (Figure 3H). The embryonic flower phenotype of *icu11-5 cp2-1* and *emf2-3* is therefore likely to be a direct consequence of the misregulation of genes composing clusters 1, 5, and 7. Cluster 6 comprised 140 genes that are downregulated in Col-0 and *cp2-1* seedlings, moderately downregulated in *icu11-5* seedlings, and upregulated in *icu11-5 cp2-1* and *emf2-3* embryonic flowers, as well as in Col-0 inflorescences (Figure 3H).

GO and protein domain enrichment analyses of each k-cluster revealed that the categories enriched in cluster 5 are related to responses to different stimuli, regulation of metabolic processes, and regulation of transcription, being mainly represented by WRKY, NAC, and the Ethylene Responsive Factor (ERF) transcription factors. In cluster 6, only the positive regulation of transcription mediated by RNA polymerase II category was enriched, represented by 10 MADS-box genes (Supplemental Data Set 8B). In cluster 1, we observed enrichment in processes related to photosynthesis, while cluster 7 included more enriched categories related to responses to biotic and abiotic stresses. In conclusion, our RNA-seq analyses provide evidence of the substantial alteration of transcript levels in the *icu11-5 cp2-1* double mutant compared to the profiles of the *icu11-5* and *cp2-1* single mutants. Additionally, the transcriptomic profile of *icu11-5 cp2-1* resembles that of mutants affected in genes encoding the PRC2 core components and accessory proteins, in particular the *emf2-3* mutant. Finally, some of the misregulated genes (clusters 1 and 7) in the *icu11-5 cp2-1* embryonic flowers are expressed as they are in the wild-type reproductive organs of Col-0 inflorescences.

### Genes misregulated in the *icu11-5 cp2-1* double mutant are enriched in PRC2 targets and genes marked with H3K27me3

A previous report suggested that ICU11 is a H3K36me3 demethylase, based on the substantial decrease of the H3K27me3 repressive mark seen in the *icu11-3* mutant (Bloomer et al., 2020). We performed a comparative analysis of the genes misregulated in *icu11-5, icu11-5 cp2-1*, and *emf2-3* with genes known to be marked by H3K27me3, H2AK121ub, and H3K36me3 in Col-0 by chromatin immunoprecipitation (ChIP)-seq published data, which are deposited by PRC2, PRC1 and SET DOMAIN-CONTAINING GROUP 8 (SDG8), respectively (Li et al., 2015; Merini et al., 2017; Sanders et al., 2017; Yang et al., 2014; Zhou et al., 2017). We determined that the genes marked with H3K27me3 and H2AK121ub in Col-0 are significantly overrepresented among the DEGs of *icu11-5, icu11-5 cp2-1*, and *emf2-3*; genes marked with H3K36me3 in Col-0 were underrepresented among the DEGs of these mutants (Figure 4A and Supplemental Table S3). Genes individually targeted by the TRB1, EMF1, and LHP1 accessory proteins of PRC2 and by both CLF and SWN (Kim et al., 2012; Shu et al., 2019; Veluchamy et al., 2016; Zhou et al., 2018) were significantly enriched among the genes misregulated in *icu11-5, icu11-5 cp2-1,* and *emf2-3*. The TRB1 targets, however, were underrepresented among the downregulated genes in *icu11-5* and *icu11-5 cp2-1*. Taken together, these data suggest that most genes misregulated in the *icu11-5 cp2-1* double mutant are direct targets of PRC2 or its accessory proteins.

**Figure 4.**
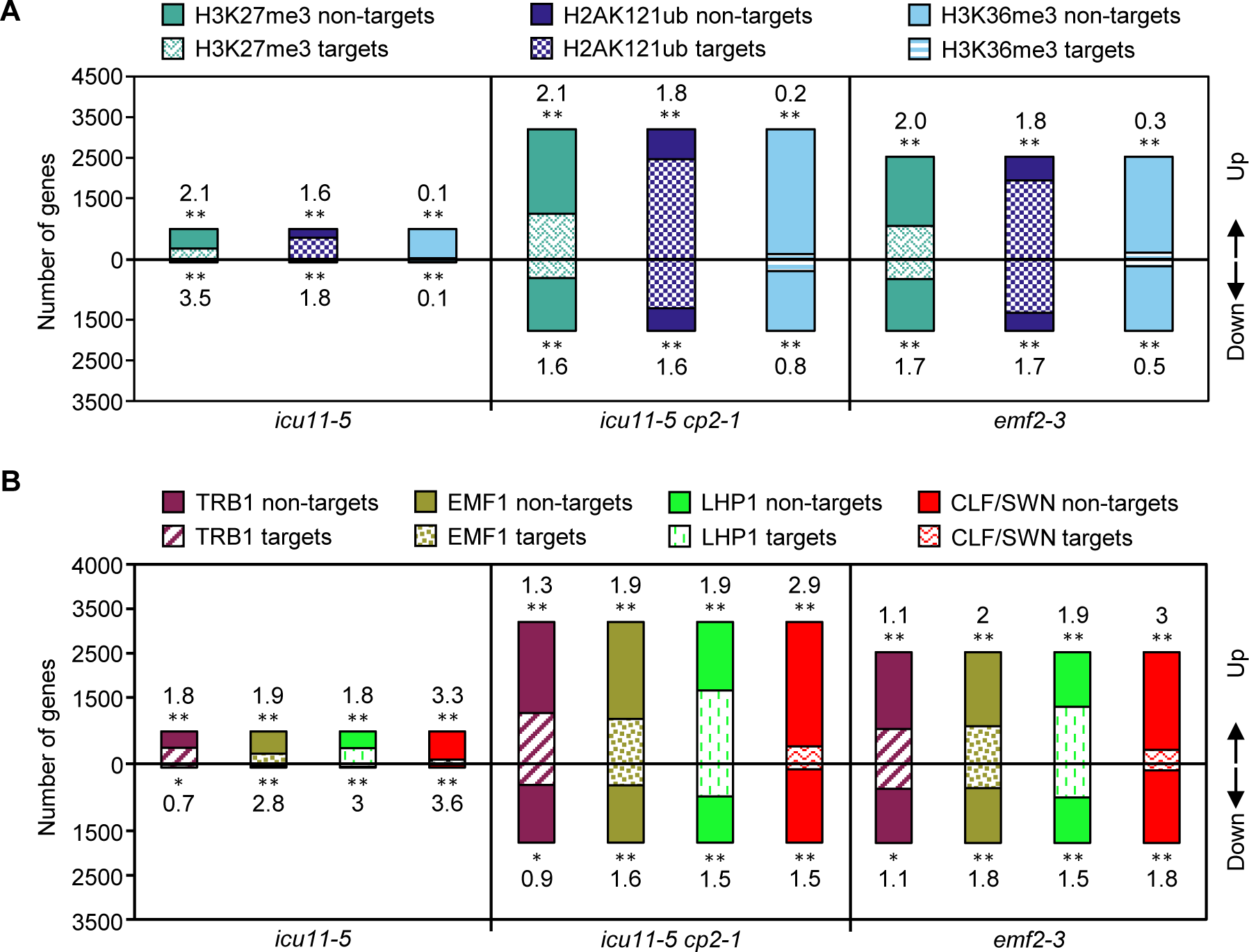
Integrated comparison of the chromatin immunoprecipitation-seq data and transcriptomic profiles in *icu11-5, icu11-5 cp2-1,* and *emf2-3* and lists of genome-wide histone mark distributions or protein targets in Col-0. (A) Overlapping fraction of upregulated and downregulated genes in the indicated mutants with genes marked by H3K27me3, H2AK121ub, and H36K36me3 in Col-0. (B) Overlapping fraction of upregulated and downregulated genes in the indicated mutants with genes bound by the TRB1, EMF1, LHP1, CLF, and SWN proteins in Col-0. Numbers indicate the enrichment factor of overlapping fractions [(number of common genes **×** number of total Arabidopsis genes)/(number of genes in list 1 **×** number of genes in list 2)], where enrichment factors > 1 or < 1 indicate more or less overlap than expected between the two independent gene lists, respectively. Asterisks indicate a significant overlap between the RNA-seq and ChIP-seq lists in a Fisher’s exact test (\**P* < 0.05 and \*\**P* < 0.01).

## DISCUSSION

### The TAP- and BiFC-based protein-protein interaction profiles of ICU11 and CP2 partially overlap, pointing to their roles as PRC2 accessory proteins

We previously showed that *ICU11* and *CP2* are close paralogs whose encoded proteins behave as components of the epigenetic machinery of Arabidopsis, which display unequal functional redundancy (Mateo-Bonmatí et al., 2018; Nadi et al., 2023). A co-IP analysis indicated that ICU11 is a PRC2 accessory protein (Bloomer et al., 2020). Very recently, both ICU11 and CP2 were shown by co-IP followed by mass spectrometry to interact with the PRC2 accessory proteins TRB1, TRB2 and TRB3, although this result was not discussed by the authors (Wang et al., 2023).

Here, we aimed to define the ICU11 and CP2 interactomes and their potential overlap in order to ascertain their epigenetic activities. We confirmed the physical interactions between ICU11 and the core components and accessory proteins of PRC2, through experimental approaches that are complementary to co-IP: TAP-based screens, and heterologous BiFC-based assays. Through these techniques, we also provide evidence that CP2 is likely to play a role as a PRC2 accessory protein, as ICU11 appears to do.

Our TAP-based screens revealed different protein-protein interaction profiles for ICU11 and CP2, despite their unequal functional redundancy; however, in our BiFC assays, ICU11 and CP2 showed similar *in vivo* interaction profiles. Other examples of partially or completely divergent results obtained from different methods of studying protein-protein interactions have been published for Arabidopsis. One of these examples is given by the pentatricopeptide repeat proteins SLOW GROWTH 2 (SLO2) and MITOCHONDRIAL EDITING FACTOR 57 (MEF57), which appeared to interact in mitochondria based on a BiFC assay, but did not interact using co-IP assays (Andrés-Colás et al., 2017). The Arabidopsis circadian clock regulators SPINDLY (SPY) and PSEUDO-RESPONSE REGULATOR 5 (PRR5) interacted in co-IP followed by mass spectrometry, as well as in co-IP followed by the identification of interactors by Western Blot and BiFC assays, but not in yeast two-hybrid (Y2H) assays; in addition interaction between SPY and GIGANTEA (GI) was detected using Y2H but not by co-IP either followed by mass spectrometry or Western Blot (Wang et al., 2020).

Our results indicate that CP2 can bind to TRBs and other proteins related to PRC2, and suggest that CP2 has the potential to bind to ICU11 interactors with less affinity than ICU11. A similar observation has been made in budding yeast (*Saccharomyces cerevisiae*), in a protein fragment complementation assay that was performed for 56 pairs of redundant paralogs. For 22 such pairs, one paralog had weaker detectable interactions than the other because of lower abundance or affinity; when the latter was lost, the former compensated for its function by binding to the same partners (Diss et al., 2017). It is of note that for compensating pairs, there was no detectable change in the level of expression of the functional paralog when the other was deleted. The same appears to hold for *CP2* in an *icu11* background, as *CP2* is not upregulated in the *icu11-5* mutant (this work) or in *icu11-1* (Mateo-Bonmatí et al., 2018). Another example is provided in human T cells by the retinoblastoma-associated protein p130, which binds to EARLY 2 FACTOR (E2F) transcription factors to control cell proliferation by gene repression. When p130 is depleted, its paralogous p107 gains new interactions with E2F proteins to compensate for the absence of p130 (Mulligan et al., 1998).

### In addition to their morphological phenotypes, the molecular phenotypes of the *icu11-5 cp2-1* and *emf2-3* embryonic flowers are similar

Our RNA-seq analyses revealed that about 21% of Arabidopsis genes were significantly misregulated in the *icu11-5 cp2-1* lethal embryonic flowers. Alongside its synergistic morphological phenotype, the *icu11-5 cp2-1* double mutant also had six times more misregulated genes than the *icu11-5* single mutant, overlapping to a large extent with misregulated genes in mutants affected in the PcG genes with strongly aberrant phenotypes, such as the *emf2-3* embryonic flowers and the *clf-29 swn-21* callus-like seedlings (Wang et al., 2016). Like in the *emf1* and *emf2* single mutants, in which many genes related to photosynthesis are repressed (Moon et al., 2003), ICU11 and CP2 appear to be involved in the positive regulation of photosynthesis, photosystem II assembly, the response to light stimulus, and auxin biosynthesis and signaling. Except for the latter, the downregulation of these genes in *icu11-5 cp2-1* and *emf2-3* embryonic flowers is also shown in wild-type Col-0 inflorescences (Kim et al., 2010; Moon et al., 2003).

During vegetative growth, ICU11 and/or CP2 seem to negatively regulate hundreds of genes to ensure the proper repression of genes that induce flowering, the formation of flower organs, the response to phytohormones, and abiotic stress. Genes encoding homeobox, MADS-box, and MYB, AP2/ERF and NAM/NAC domain transcription factors were enriched among the *icu11-5 cp2-1* upregulated DEGs. These genes were also highly expressed in Col-0 inflorescence meristems, where they play a crucial role in floral meristem development (Jofuku et al., 1994; Zhang et al., 2014; Zhang et al., 2009). Our k-mean clustering analysis of the most differentially regulated genes allowed the identification of a set of 1573 genes (clusters 1, 6, and 7 in Figure 3) that are expressed similarly in the *icu11-5 cp2-1* and *emf2-3* embryonic flowers and Col-0 inflorescences. Another set encompassed 1377 genes (clusters 2, 3, and 4) that are exclusively differentially expressed in Col-0 inflorescences. Finally, a set of 610 genes (cluster 5) comprised those highly upregulated only in *icu11-5 cp2-1* and *emf2-3,* and slightly upregulated in the *icu11-5* single mutant seedlings. This last set of genes was characterized by GO enrichment related to the responses to chemicals, oxygen-containing compounds, drugs, chitin and inorganic substances and stimuli, the regulation of metabolic and biosynthetic processes, with 54 genes involved in regulation of transcription. Our results explain the embryonic flower phenotypes of *icu11-5 cp2-1* and *emf2-3*, given that there are 696 and 817 common up- and down-regulated genes with the wild-type inflorescence, respectively (clusters 1, 6, and 7), but also their failure to form a proper inflorescence, as expected from the 1377 genes that behave differently in the Col-0 inflorescences (clusters 2, 3, and 5). In conclusion, the similarity of not only the morphological but also the molecular phenotypes of *icu11-5 cp2-1* and *emf2-3* provides further support for the hypothesis that both ICU11 and CP2 are PRC2 accessory proteins.

### Our interactomic and transcriptomic data suggest that CP2 can replace ICU11

Bloomer et al. (2020) proposed that ICU11 is a H3K36me3 demethylase. The depletion of a protein involved in the removal of an activating mark is expected to yield predominantly upregulated genes, which is in line with the pattern of misregulation detected here. This pattern has also been observed for lack-of-function alleles of the *JMJ17* and *JMJ14* genes, whose encoded proteins remove the H3K4me1/2/3 activating marks (Huang et al., 2019; Ning et al., 2015). The H3K36me3 mark antagonizes the deposition of H3K27me3 by PRC2 (Yang et al., 2014), which may explain the requirement of ICU11 for the deposition of the latter mark at the *FLC* locus by PRC2 during vernalization (Bloomer et al., 2020). It is therefore reasonable to assume that the transcriptomic profile of *icu11-5 cp2-1* is similar to that of a strong PcG mutant, such as *emf2-3*, because PRC2 cannot deposit H3K27me3 when the H3K36me3 mark cannot be removed.

Our comparison of the transcriptional misregulation of *icu11-5, icu11-5 cp2-1* and *emf2-3* with published ChIP-seq data reveals that genes marked by H3K27me3 and H2AK121ub or targeted by PRC2 core components and accessory proteins are overrepresented among the genes misregulated in these three mutants; the morphological and transcriptomic alterations observed for these genotypes are likely to be due to defective PRC2 repression on a substantial set of genes. If ICU11 and CP2 targets are not marked with H3K36me3 in Col-0, this would explain the underrepresentation of H3K36me3 marked genes among the differentially expressed genes in *icu11-5* and *icu11-5 cp2-1*.

We propose that ICU11 interacts with PRC2 core and accessory proteins, some of which recruit ICU11 to their target genes, so ICU11 can demethylate H3K36me3 and PRC2 can deposit H3K27me3 afterwards (Figure 5A). In the *icu11* mutants, although CP2 has less affinity for PRC2 core and accessory proteins, it can substitute ICU11 and demethylate H3K36me3 (Figure 5B). In the *icu11-5 cp2-1* double mutant, the H3K36me3 mark cannot be removed, leading to an impairment of PRC2 repression, resulting in the characteristic embryonic flower phenotype (Figure 5C). Since this does not explain the wild-type function of CP2, further research will be required to assess the specific function of CP2.

**Figure 5.**
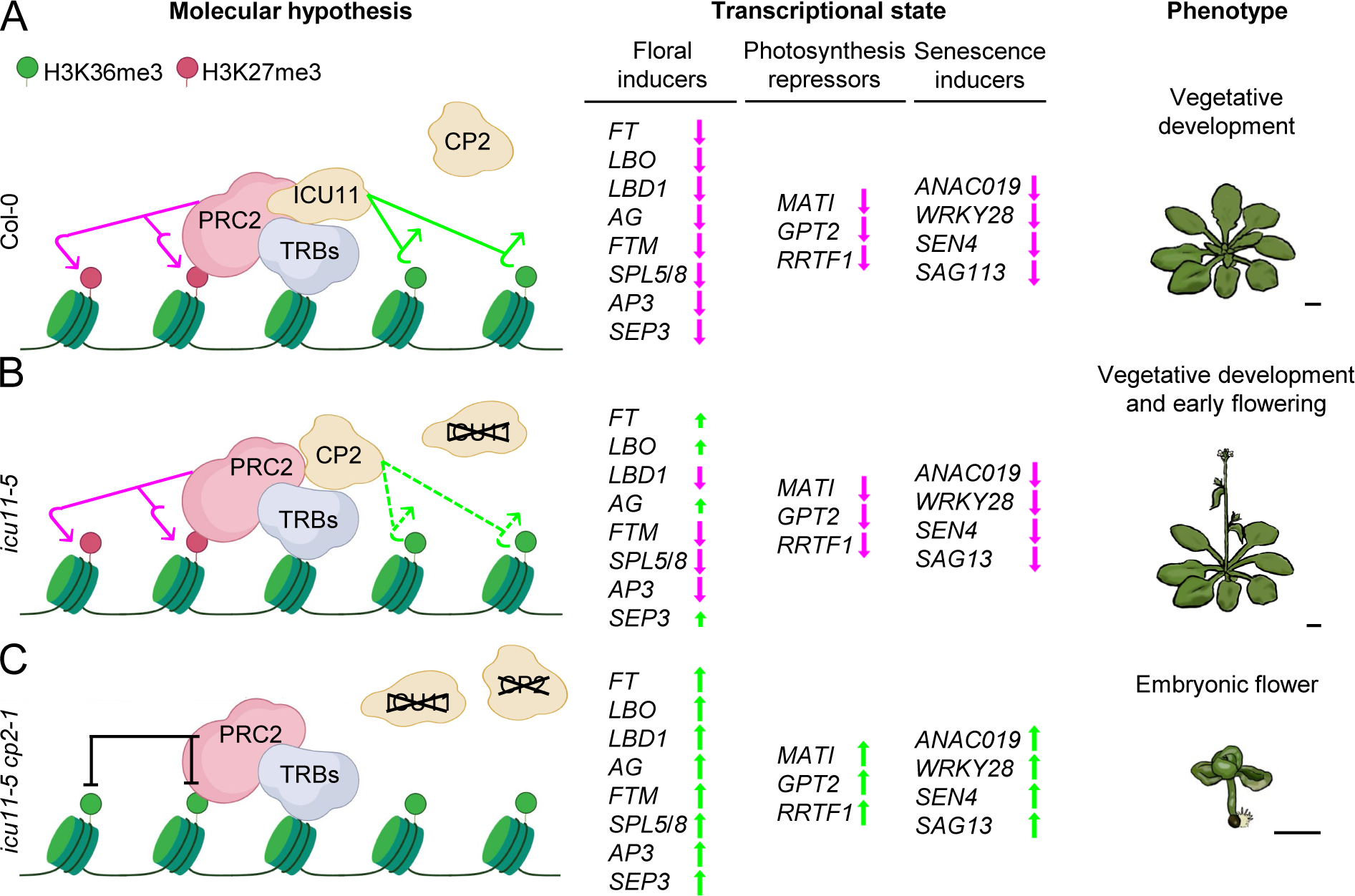
Model of the molecular role of ICU11 and CP2 and the effects of their depletion on transcription and phenotype in Arabidopsis. (A) In the wild-type Col-0, ICU11 may bind to the TRB1, TRB2, and TRB3 accessory proteins of PRC2, which recruit ICU11 to its target loci to remove the H3K36me3 activation mark. This enables PRC2 to deposit the repressive mark H3K27me3, leading to the repression of flower development genes, senescence inducers, and photosynthesis repressor genes, promoting proper vegetative development. (B) In the *icu11-5* mutant, CP2 can only partially compensate for the absence of ICU11 because of its lower affinity for TRB proteins and its less efficient removal of H3K36me3, decreasing the PRC2 repressive capacity, which results in an early flowering phenotype. (C) In the *icu11-5 cp2-1* double mutant, the presence of H3K36me3 at the target loci of ICU11 and CP2 impedes the deposition of H3K27me3, leading to an upregulation of floral and senescence inducers and photosynthetic repressors, resulting in the embryonic flower phenotype. **∠**: full or partial depletion of a protein. ⤤: H3K36me3 removal. ⤥: H3K27me3 deposition. ┴: inhibition of PRC2-mediated H3K27me3 deposition. ⬆: transcriptional activation. ⬇: transcriptional repression. Scale bars indicate 2 mm.

### TRB and NAC proteins may recruit both ICU11 and CP2

ICU11 was previously described as a putative H3K36me3 demethylase, required for the removal of H3K36me3 during vernalization to allow the deposition of H3K27me3 (Bloomer et al., 2020). The interaction of both ICU11 and CP2 with TRB1, TRB2, and TRB3 suggests that these TRB proteins may recruit ICU11 and CP2 to their chromatin targets (Zhou et al., 2018). This hypothesis is reinforced by the similar embryonic flower phenotypes and transcriptomic profiles of the *icu11 cp2* double mutants and the *trb1-2 trb2-1 trb3-2* triple mutant.

The Arabidopsis genome encodes 21 2OGD proteins of the JMJ class, some of which are known to be involved in histone demethylation (Lu et al., 2008; Nadi et al., 2018). The H3K4 demethylase JMJ14 interacts with the NAC050 and NAC052 transcription factors through its phenylalanine/tyrosine-rich C-terminal (FYRC) domain, and plays an essential role in controlling flowering time (Ning et al., 2015). Here, we showed that ICU11 and CP2 also interact with NAC050 and NAC052, even though unlike JMJ14, ICU11 and CP2 do not have FYRC or FYRN domains. It is of note that neither NAC050 nor NAC052 were identified as ICU11 interactors by Bloomer et al. (2020), although JMJ14 was. NAC050 and/or NAC052 might recruit ICU11 and CP2 to their targets.

We also found three paralogous nuclear proteins that were not previously described as ICU11 interactors: At5g66000, At3g17460 and AT4G35510. At3g17460 and At4g35510 have a PHD domain. Bloomer et al. (2020) showed that the protein encoded by At5g66000 interacts with ICU11, EMF1 and CLF, that At3g17460 interacts with ICU11 and that At4g335510 interacts with CLF. We also showed that CP2 interacts with DRMY1 and DP1. Given that both ICU11 and CP2 lack any known DNA- or chromatin-binding domain, our results indicate they interact with proteins that may mediate their interaction with DNA or chromatin. Taken together, our interactome and transcriptome data confirm that ICU11 is a PRC2 accessory protein, and strongly suggest that CP2 also does this role for the correct deposition of H3K27me3 by PRC2.

## METHODS

### Plant materials, culture conditions, and crosses

The Nottingham Arabidopsis Stock Center (NASC) provided seeds for the wild-type *Arabidopsis thaliana* (L.) Heynh. accession Columbia-0 (Col-0, N1092), and the mutants *cp2-1* (N861581, in the Col-0 genetic background) and *emf2-3* (N16240, in Col-0). The *icu11-5* (in Col-0) single mutant was obtained using CRISPR/Cas9 mutagenesis and was described previously by Nadi et al. (2023). The presence and position of all mutations were confirmed by PCR amplification using gene-specific primers and, if required, Sanger sequencing (Supplemental Table S4).

Unless otherwise stated, plants were grown under sterile conditions in 150-mm Petri plates containing 100 ml half-strength Murashige and Skoog (MS) agar medium with 1% (w/v) sucrose at 20°C ± 1°C, 60–70% relative humidity, and continuous illumination at ∼75 µmol m^– 2^ s^–1^, as previously described (Ponce et al., 1998). The crosses were performed as previously described (Quesada et al., 2000). Unless otherwise stated, all plants studied in this work were homozygous for the indicated mutations.

### Gene constructs

All inserts were PCR amplified using Phusion High Fidelity Polymerase (Thermo Fisher Scientific, Waltham, MA, USA), primers containing *att*B sites at their 5′ ends (Supplemental Table S4), and Col-0 complementary DNA (cDNA) as a template. The PCR products were purified using an Illustra GFX PCR and Gel Band Purification Kit (Cytiva, Marlborough, MA, USA) and then cloned into the pGEM-T Easy221 vector and transferred to *Escherichia coli* DH5α cells, as previously described (Mateo-Bonmatí et al., 2018).

### Tandem affinity purification assays

To obtain the GSRhino-TAP-tagged ICU11 or CP2 fusions (Supplemental Table S5), the pGEM-T Easy221 vector harboring the *ICU11* or *CP2* full-length coding sequences without their stop codons, together with the vectors pEN-L4-2-R1 and pEN-R2-GSrhinotag-L3, were recombined into the pKCTAP destination vector, as previously described (Van Leene et al., 2015). PSB-D Arabidopsis cell suspension cultures were transformed with *Agrobacterium tumefaciens* cells carrying the constructs and the TAP purification of the GSRhino-TAP-tagged ICU11 and CP2 fusions was performed as previously described (García-León et al., 2018; Van Leene et al., 2015). Two independent TAP assays were performed for each fusion protein. Proteins were identified using nano liquid chromatography–mass spectrometry (LC–MS)/MS at the Centro Nacional de Biotecnología (CNB, Madrid). Tandem mass spectra were searched against the Araport11 annotation of the Arabidopsis genome (Cheng et al., 2017) using the MASCOT search engine (Perkins et al., 1999). Experimental background proteins were subtracted based on 40 TAP experiments performed on wild-type cultures and cultures accumulating GSRhino-TAP-tagged GUS, RFP, and GFP fusion proteins (Van Leene et al., 2010).

### BiFC assays in *Nicotiana benthamiana* leaves and confocal microscopy

To obtain translational fusions for the BiFC assays, the pGEM-T Easy221 vector harboring the full-length *ICU11*, *CP2*, *TRB1*, *TRB2*, *TRB3*, *CLF*, *LHP1*, and *SWN* coding sequences, including their stop codons, were individually recombined with the pSITE-nEYFP-C1 or pSITE-cEYFP-C1 vectors (Martin et al., 2009). The nEYFP-UBP12 and nEYFP-UBP13 constructs were kindly provided by Dr. Claudia Köhler (Max Planck Institute, Postdam, Germany) Derkacheva et al., 2016). The BiFC constructs (Supplemental Table S5) were transformed into Agrobacterium (*Agrobacterium tumefaciens*) strain GV3101 (C58C1 Rif^R^) cells, which were the grown in suspension as previously described (Derkacheva et al., 2016; Goodin et al., 2002). Briefly, the cells were grown overnight and resuspended in infiltration medium (10 mM MgCl_2_, 150 μg/ml acetosyringone, and 10 mM MES-KOH, pH 5.6) to a final optimal density (OD_600_) ≤ 1. After 3 h at room temperature, the Agrobacterium cell suspension was used to infiltrate the leaf abaxial surface of three-to five-week-old *Nicotiana benthamiana* plants. Leaf tissue samples were water-mounted for confocal visualization 48 h after infiltration.

Confocal microscopy was performed with a Nikon D-Eclipse C1 confocal microscope equipped with a Nikon DS-Ri1 camera and processed with the operator software EZ-C1 (Nikon, Tokyo, Japan). YFP was excited at 488 nm with an argon ion laser, and the emission signal was collected between 520 nm and 582 nm. The nuclei of the infiltrated leaves were stained with a 0.2 μg ml^−1^ 4′,6-diamidino-2-phenylindole (DAPI) solution (Sony Biotechnology, San José, CA, USA). DAPI was excited at 408 nm with a diode laser, and detected with a 450/35 nm filter.

### RNA-seq analyses

Total RNA was isolated from 100 mg of pooled aerial tissues from Col-0, *icu11-5, cp2-1, icu11-5 cp2-1,* or *emf2-3* seedlings, collected 10 das, and Col-0 inflorescences, collected 40 das, using TRIzol (Thermo Fisher Scientific). The RNA quality of the samples was checked with a 2100 Bioanalyzer (Agilent Technologies, Santa Clara, CA, USA), and its RNA integrity number (RIN) was always ≥ 6.8. More than 10 µg of RNA per sample was sent to Novogene (Cambridge, UK) for library preparation and massive sequencing on an Illumina Novaseq 6000 (Illumina, San Diego, CA, USA).

Raw reads were pre-processed using fastp (v.0.21.0; Chen et al., 2018) with default parameters for read trimming, adapter removal, and low-quality read filtering. Pre-processed reads were then aligned to the TAIR10 reference genome (Lamesch et al., 2012) using HISAT2 (v.2.2.0; Kim et al., 2019), with argument “—dta-cufflinks” for downstream compatibility (Supplemental Table S6). Cufflinks (v.2.2.1; Trapnell et al., 2012) was then used for transcript assembly using the TAIR10 structural annotation for reference, and transcripts were quantified with htseq-count to generate read count files (v.0.11.5; Anders et al., 2015). Read counts were normalized with DESeq2 (v.1.30.0; Love et al., 2014), which was then used to detect DEGs) between the sample and control pairs using the combined criteria |log2-fold-change| > 1 and *p*.adj value < 0.05. Volcano plots were obtained with the volcano plot tool of Galaxy (www.usegalaxy.org; The Galaxy Community, 2022). For principal component analysis we used NetworAnalyst tool (Zhou et al., 2019).

Both GO and Protein Domain enrichment analyses of the DEGs were performed using DAVID Bioinformatics tool (v.6.8; Huang da et al., 2009) with default parameters. Heatmaps were obtained using the heatmap.2 function from the gplots R package (v.20 3.0.1) using a total of 11116 genes that were misregulated in at least one of the genotypes under study. Additional RNA-seq data were downloaded from the Gene Expression Omnibus (https://www.ncbi.nlm.nih.gov/geo/) under accession number SRP056594 and from the European Nucleotide Archive (https://www.ebi.ac.uk/ena/browser/) under accession numbers ERP022017 and ERP009986. A principal component analysis and k-means clustering was performed using normalized counts on the iDEP 9.1 web 133 application (Ge et al., 2018). ChIP-seq data for cross analysis with RNA-seq were obtained from work published by Kim et al. (2012), Li et al. (2015), Merini and Calonje (2015), Sanders et al. (2017), Shu et al. (2019), Veluchamy et al. (2016), Zhou et al. (2017), and Zhou et al. (2018).

### RNA isolation, cDNA synthesis, and qPCR

For the RT-qPCR, three biological replicates of seedling aerial tissues were collected 10 das and immediately frozen in liquid nitrogen. Total RNA was extracted using TRIzol (Thermo Fisher Scientific). The removal of contaminating DNA, cDNA synthesis, and qPCR were performed as previously described (Mateo-Bonmatí et al., 2018). Each reaction was performed in triplicate and the relative quantification of gene expression was performed using the 2^–ΔΔC^_T_ method (Livak and Schmittgen, 2001; Schmittgen and Livak, 2008) with the *ACTIN2* gene (At3g18780) as a control. All PCR reactions were performed on an Applied Biosystems Step One Plus System (Thermo Fisher Scientific). All PCR primers are listed in Supplemental Table S4; for the mean ΔCT statistical comparisons, a Mann-Whitney *U* test was performed.

### Accession numbers

Sequence data from this article can be found at The Arabidopsis Information Resource (http://www.arabidopsis.org) under the following accession numbers: *ICU11* (At1g22950), *CP2* (At3g18210), *EMF2* (AT5G51230), *SWN* (AT4g02020)*, CLF* (AT2g23380), *TFL2/LHP1* (At5g17690), *AG* (At4g18960), *SHP2* (AT2g42830), *STK* (AT4g09960), *EARLI1*(AT4g12480), *PRN1* (AT3g59220), *RNS1* (AT2g02990), *TRB1* (AT1g49950)*, TRB2* (AT5g67580), *TRB3* (AT3g49850), *TRB4* (AT1g17520), and *TRB5* (AT1g72740). The raw RNA-seq data were deposited in the Sequence Read Archive (SRA, https://www.ncbi.nlm.nih.gov/sra) database under the following accession number: PRJNA1081349.

## AUTHOR CONTRIBUTIONS

J.L.M. conceived and supervised the study, provided resources, and obtained funding. R.N., L.J.-V., and J.L.M. designed the methodology. R.N., L.J.-V., S.D.L, Y.F., and V.R. performed the research. R.N., L.J.-V. and J.L.M. wrote the original draft. All authors reviewed and edited the manuscript.

## ACKNOWLEDGMENTS

The authors wish to thank C. Köhler and I. Fudal for providing constructs, M.R. Ponce and E. Mateo-Bonmatí for critical reading of the manuscript, J.M. Serrano and J. Castelló for their excellent technical assistance, and S. Vivo and C. Torralbo for their help in BiFC assays. Research in the laboratory of J.L.M. was supported by grants from the Ministerio de Ciencia e Innovación of Spain (PGC2018-093445-B-I00 and PID2021-127725NB-I00 [MCI/AEI/FEDER, UE]) and the Generalitat Valenciana (CIPROM/2022/2). R.N. and L.J.-V. held predoctoral fellowships from the Generalitat Valenciana (GRISOLIAP/2016/131) and the Ministerio de Universidades of Spain (FPU16/03772), respectively.

## COMPETING INTERESTS

The authors declare no competing interests.

## Supplemental material

Supplemental Figure S1. Diagram of the protein-protein interactions of ICU11 and CP2 detected in tandem affinity purification (TAP)-based screens.

Supplemental Figure S2. Peptides from TRB proteins identified using Liquid Chromatography Electrospray Ionization and Tandem Mass Spectrometry (LC-ESI-MS/MS) in ICU11 and CP2 TAP-based screens.

Supplemental Figure S3. Controls used for the Bimolecular Fluorescence Complementation (BiFC) assays.

Supplemental Figure S4. Validation by reverse transcription-quantitative PCR (RT-qPCR) of some of the genes found to be upregulated in our RNA-seq analyses.

Supplemental Table S1. Selected ICU11 and CP2 interactors identified in TAP-based screens.

Supplemental Table S2. Number of differentially expressed genes in *cp2-1* and *icu11-5* seedlings, *icu11-5 cp2-1* and *emf2-3* embryonic flowers, and Col-0 inflorescences, compared to the Col-0 seedlings.

Supplemental Table S3. Enrichment of overlapping fractions of chromatin immunoprecipitation (ChIP)-seq and transcriptomic profiles and their statistical significance.

Supplemental Table S4. Primer sets used in this work.

Supplemental Table S5. TAP and BiFC constructs.

Supplemental Table S6. Quality control summary of the RNA-seq analyses

Supplemental Data Set DS1. Protein identification in ICU11 and CP2 TAP-based screens.

Supplemental Data Set DS2. Differentially expressed genes in the RNA-seq analyses of *icu11-5* and *cp2-1* seedlings, *icu11-5 cp2-1* and *emf2-3* embryonic flowers, and Col-0 inflorescences.

Supplemental Data Set DS3. Protein domains and biological process gene ontology terms enriched among genes deregulated in *icu11-5* seedlings.

Supplemental Data Set DS4. Protein domains and biological process gene ontology terms enriched among genes deregulated in *cp2-1* seedlings.

Supplemental Data Set DS5. Protein domains and biological process gene ontology terms enriched among genes deregulated in *icu11-5 cp2-1* embryonic flowers.

Supplemental Data Set DS6. Protein domains and biological process gene ontology terms enriched among genes deregulated in *emf2-3* embryonic flowers.

Supplemental Data Set DS7. Protein domains and biological process gene ontology terms enriched among genes deregulated in Col-0 inflorescences.

Supplemental Data Set DS8. Biological process gene ontology enrichment analysis from k-means gene clustering.

## Notes

### Competing Interest Statement

The authors have declared no competing interest.

