## Supplemental Material for "Overlapping roles of Arabidopsis INCURVATA11 and CUPULIFORMIS2 as Polycomb Repressive Complex 2 accessory proteins"

[1]Instituto de Bioingeniería, Universidad Miguel Hernández, Campus de Elche, 03202 Elche, Spain; [2]Centro Nacional de Biotecnología, CNB-CSIC, Madrid, Spain. *These authors contributed equally to this work.

### Supplemental Figures and Tables

Supplemental Material included in this file:

- Supplemental Figures S1-S4
- Supplemental Tables S1-S6

Supplemental Material not included in this file:

- Supplemental Datasets DS1-DS8

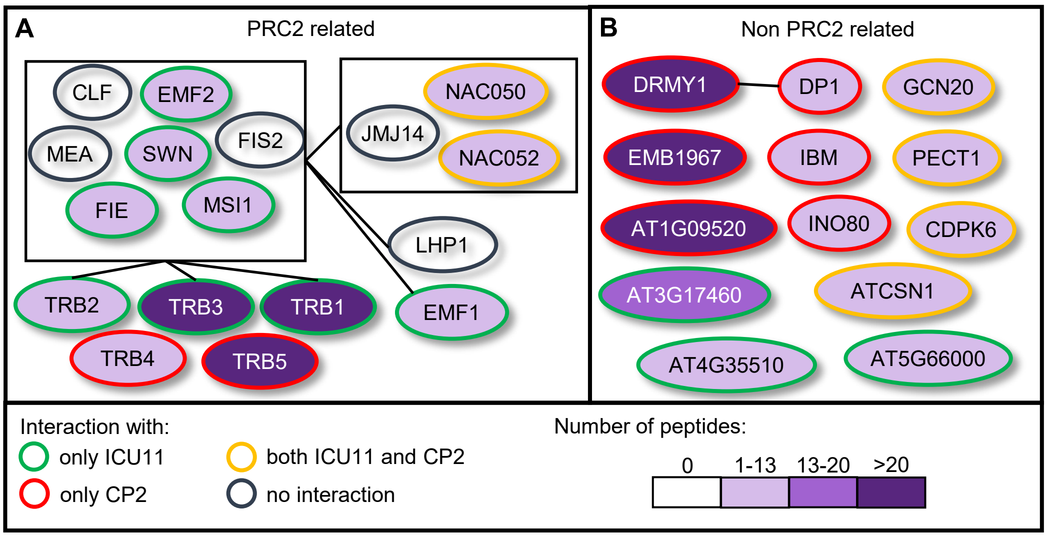


**Supplemental Figure S1.** Diagram of the protein-protein interactions of ICU11 and CP2 detected in Tandem Affinity Purification (TAP)-based screens. Proteins that interacted with ICU11 and CP2 in TAP assays, classified as (A) known to be related to the PRC2, or (B) not related. Proteins are represented as ellipses outlined in green, red, yellow, and black lines, depending upon their interaction with only ICU11, only CP2, both ICU11 and CP2, or no interaction with ICU11 or CP2, respectively. Black lines indicate previously known interactions. The violet color intensity scale represents the maximum number of peptides from each protein identified as detailed in Supplemental Table S1. Boxes represent the previously described complexes or groups of interacting proteins, and include some that we found not to interact with ICU11 or CP2 (the CLF, MEA and FIS2 core components of PRC2, and the JMJ14 accessory protein).

AT1G49950 (TRB1; 53.33%)

MGAPKQKWTQEEESALKSGVIKHGPGKWRTILKDPEFSGVLYLRSNVDLKDKWRNMSVMANGWGSREKSRLAVKRTFSL
PKQEENSLALTNSLQSDEENVDATSGLQVSSNPPPRRPNVRLDSLIMEAIATLKEPGGCNKTTIGAYIEDQYHAPPDFK
RLLSTKLKYLTSCGKLVKVKRKYRIPNSTPLSSHRRKGLGVFGGKQRTSSLPSPKTDIDEVNFQTRSQIDTEIARMKSM
NVHEAAAVAAQAVAEAEAAMAEAEEAAKEAEAAEAEAEAAQAFAEEASKTLKGRNICKMMIRA

AT5G67580 (TRB2; 42.14%)

MGAPKQKWTPEEEAALKAGVLKHGTGKWRTILSDTEFSLILKSRSNVDLKDKWRNISVTALWGSRKKAKLALKRTPPGT
KQDDNNTALTIVALTNDDERAKPTSPGGSGGGSPRTCASKRSITSLDKIIFEAITNLRELRGSDRTSIFLYIEENFKTP
PNMKRHVAVRLKHLSSNGTLVKIKHKYRFSSNFIPAGARQKAPQLFLEGNNKKDPTKPEENGANSLTKFRVDGELYMIK
GMTAQEAAEAAARAVAEAEFAITEAEQAAKEAERAEAEAEAAQIFAKAAMKALKFRIRNHPW

AT3G49850 (TRB3; 50.51%)

MGAPKLKWTPEEETALKAGVLKHGTGKWRTILSDPVYSTILKSRSNVDLKDKWRNISVTALWGSRKKAKLALKRTPLSG
SRQDDNATAITIVSLANGDVGGQQIDAPSPPAGSCEPPRPSTSVDKIILEAITSLKRPFGPDGKSILMYIEENFKMQPD
MKRLVTSRLKYLTNVGTLVKKKHKYRISQNYMAEGEGQRSPQLLLEGNKENTPKPEENGVKNLTKSQVGGEVMIMGMTE
KEAAAAAARAVAEAEFAMAEAEEAAREADKAEAEAEAAHIFAKAAMKAVKYRMHSQTR

AT1G17520 (TRB4; 30.74%)

MGNQKLKWTAEEEEALLAGVRKHGPGKWKNILRDPELAEQLSSRSNIDLKDKWRNLSVAPGIQGSKDKIRTPKIKAAAF
HLAAAAAAAIVTPTHSGHSSPVATLPRSGSSDLSIDDSFNIVVDPKNAPRYDGMIFEALSNLTDANGSDVSAIFNFIEQ
RQEVPPNFRRMLSSRLRRLAAQGKLEKVSHLKSTQNFYKMNDNSLVQRTPHVARPKESNTKSRQQTNSQGPSISQQIVE
ASITAAYKLVEVENKLDVSKGAAEEIERLMKLAEEADEMLVIAREMHEECSQGKIMYLN

AT1G72740 (TRB5; 41.81%)

MGNQKLKWTAEEEEALLAGIRKHGPGKWKNILRDPEFADQLIHRSNIDLKDKWRNLSVPPGTQSLTNKARPAKVKEEGD
TPAADANDAVTIPRPIPTIPPPPGRRTLPSELIPDENTKNAPRYDGVIFEALSALADGNGSDVSSIYHFIEPRHEVPPN
FRRILSTRLRRLAAQSKLEKVSTFKSIQNFYKIPDPSGTKIGVPKPKETHTKLRQANNQTSADSQQMIEEAAITAACKV
VEAENKIDVAKLAAEEFEKMTKIAEENRKLLVIATEMHELCSCGETMLLA

**Supplemental Figure S2.** Peptides from TRB proteins identified using Liquid Chromatography Electrospray Ionization and Tandem Mass Spectrometry (LC-ESI-MS/MS) in ICU11 and CP2 TAP-based screens. Arabidopsis Genome Initiative (AGI) gene identifiers (AtNgNNNNN) are shown, together with the corresponding protein name and peptide coverage (in percentage) for each protein. Full-length protein sequences were obtained from The Arabidopsis Information Resource (TAIR; https://www.arabidopsis.org/). All the peptides identified were protein-specific and are shaded in black; the only exception was the peptide shaded in red, which is identical in TRB2 and TRB3. Peptide coverage is shown as a percentage, and was calculated by dividing the total number of residues of each protein by those of the peptides.

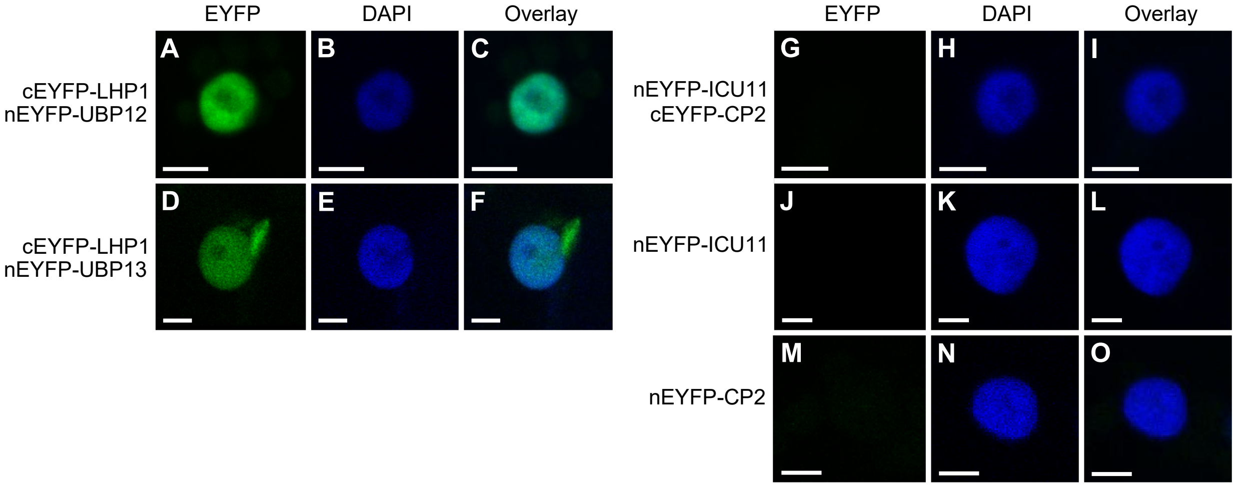


**Supplemental Figure S3.** Controls used for the Bimolecular Fuorescence Complementation (BiFC) assays. (A–F) Nuclei of *Nicotiana benthamiana* leaf cells showing BiFC of *nEYFP-UBP12* and *nEYFP-UBP13* with *cEYFP-LHP1*, which were used as positive controls. (G–O) Nuclei of *Nicotiana benthamiana* leaf cells showing no BiFC after (G–I) co-infiltration with both *nEYFP-ICU11* and *nEYFP-CP2*, or infiltration with (J–L) only *nEYFP-ICU11* or (M–O) only *cEYFP-CP2*, which were used as negative controls. Fluorescent signals correspond to EYFP (A, D, G, J, M), the DAPI nuclear dye (B, E, H, K, N), and their overlay (C, F, I, L, O). Scale bars, 5 µm.

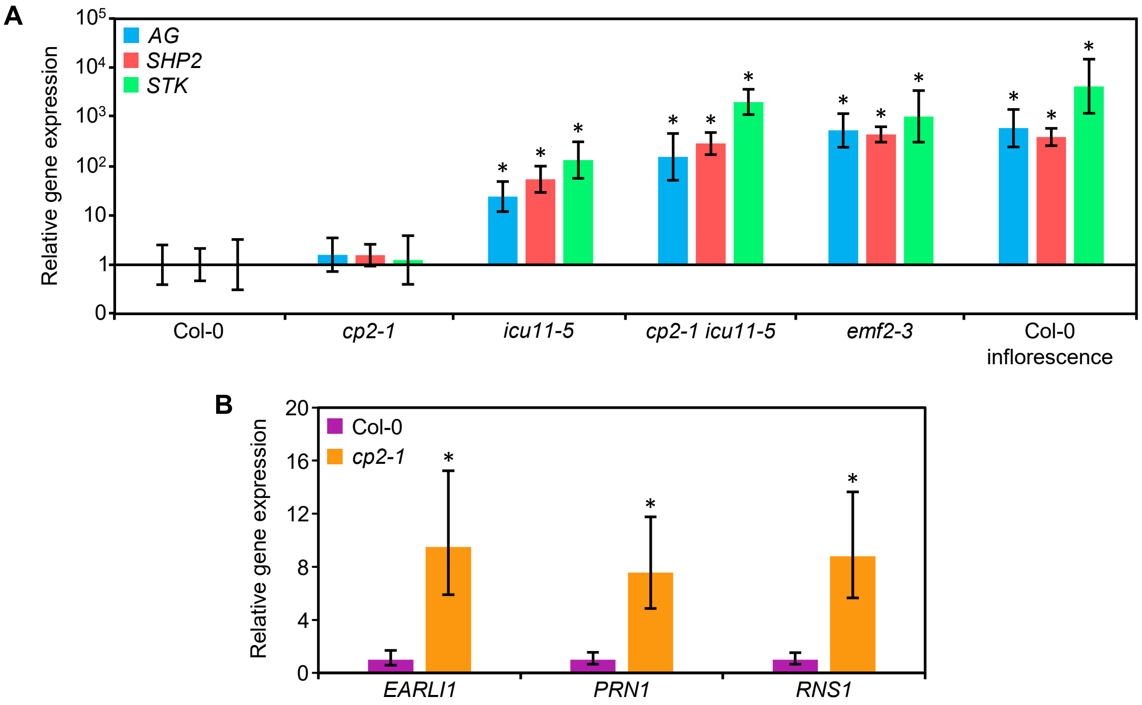


**Supplementary Figure S4.** Validation by reverse transcription-quantitative PCR (RT-qPCR) of some of the genes found to be upregulated in our RNA-seq analyses. (A) Relative expression of the *AGAMOUS* (*AG*), *SHATTERPROOF 2* (*SHP2*) and *SEEDSTICK* (*STK*) in the aerial tissues of Col-0 (only error bars are visible), *cp2-1* and *icu11-5* seedlings, and *cp2-1 icu11-5* and *emf2-3* embryonic flowers collected 10 das, as well as Col-0 inflorescences collected 40 das. (B) Relative expression of *EARLY ARABIDOPSIS ALUMINUM INDUCED 1* (*EARLI1*), *PIRIN 1* (*PRN1*) and *RIBONUCLEASE 1* (*RNS1*) in aerial tissues of Col-0 and *cp2-1* plants, collected 10 das. Values are means ± standard deviation. Asterisks indicate $2^{-\Delta\Delta C_T}$ values significantly differing from those of Col-0 in a Mann-Whitney *U* test (*$P < 0.01$; $n = 3$).

**Supplemental Table S1.** Selected ICU11 and CP2 interactors identified by TAP-based screens

| Sample | Interactor | Full name and/or annotation | Protein score |
|---|---|---|---|
| GSRhino-TAP-tagged ICU11 | TRB1[a] | PRC2 accessory protein | 962 |
| | TRB3[a] | PRC2 accessory protein | 713 |
| | AT3G17460[a] | Nuclear protein with PHD domain | 653 |
| | TRB2[a] | PRC2 accessory protein | 415 |
| | NAC052[b] | NAC DOMAIN CONTAINING PROTEIN 52; transcription factor | 208 |
| | NAC050[b] | NAC DOMAIN CONTAINING PROTEIN 50; transcription factor | 158 |
| | ALDH4[b] | ALDEHYDE DEHYDROGENASE 4 | 115 |
| | EMF1[a] | PRC2 accessory protein | 111 |
| | CDPK3[b] | CALCIUM DEPENDENT PROTEIN KINASE 6 | 133 |
| | AT4G35510 | Nuclear protein with PHD domain | 104 |
| | AT5G66000[a] | Nuclear protein with PHD domain | 92 |
| | GCN20 | GENERAL CONTROL NON-REPRESSIBLE 20 | 87 |
| | EMF2[a] | PRC2 core component | 85 |
| | PECT1 | PHOSPHORYLETHANOLAMINE CYTIDYLYLTRANSFERASE 1 | 79 |
| | FIE[a] | PRC2 core component | 71 |
| | SWN[a] | PRC2 core component | 61 |
| | CSN1 | COP9 signalosome complex subunit 1 | 65 |
| | MSI1[a] | PRC2 core component | 45 |
| GSRhino-TAP-tagged CP2 | AT1G09520 | Hypothetical nuclear protein | 1618 |
| | TRB5 | TELOMERE REPEAT BINDING FACTOR 5 | 1067 |
| | DRMY1 | DEVELOPMENT RELATED MYB-LIKE1; transcription factor | 763 |
| | EMB1967 | EMBRYO DEFECTIVE 1967 | 643 |
| | TRB4 | TELOMERE REPEAT BINDING FACTOR 4 | 374 |
| | IBM1 | IMBIBITION-INDUCIBLE 1; transcription factor | 324 |
| | RICE1 | RISC-INTERACTING CLEARING 3′-5′ EXORIBONUCLEASE 1 | 197 |
| | DGR1b | DUF642 L-GALL RESPONSIVE GENE 1 | 181 |
| | CPK3b | CALCIUM DEPENDENT PROTEIN KINASE 3 | 168 |
| | DP1 | DRMY1 PARALOG 1; transcription factor | 164 |
| | INO80 | INOSITOL REQUIRING80 | 124 |

[a]Proteins that were identified in Bloomer et al. (2020).[b]Proteins that were identified in both GSRhino-TAP-tagged ICU11 and CP2 fusions TAP assays. The Protein Score is calculated by the Mascot search engine (Perkins et al., 1999) for each protein; it is based on the probability that peptide mass matches are non-random events. If the Protein Score is equal to or greater than the Mascot Significance Level calculated for the database search, the protein match is considered to be statistically non-random at the 95% confidence interval. The Protein Score is reported as -10Log$_{10}$($P$) where $P$ is the probability that the observed match is a random event.

**Supplemental Table S2.** Number of differentially expressed genes in *cp2-1* and *icu11-5* seedlings, *icu11-5 cp2-1* and *emf2-3* embryonic flowers, and Col-0 inflorescences, compared to Col-0 seedlings

| Genes | Seedlings | | Embryonic flowers | | Col-0 inflorescence |
|---|---|---|---|---|---|
| | *cp2-1* | *icu11-5* | *icu11-5 cp2-1* | *emf2-3* | |
| Up-regulated | 23 | 738 | 3199 | 2520 | 5431 |
| Down-regulated | 5 | 78 | 1770 | 1774 | 3084 |
| Total | 28 | 816 | 4969 | 4294 | 8515 |

All samples were compared to Col-0 seedlings to detect differentially expressed genes which were filtered using padj < 0.05 and |log2FC| > 1.

**Supplemental Table S3.** Enrichment of overlapping fractions of ChIP-seq and transcriptomic profiles and their statistical significance

| Genotype | Mis-regulation | Criterion | Epigenetic marks | | | Genes bound by | | | |
|---|---|---|---|---|---|---|---|---|---|
| | | | H3K27me3 | H2AK121ub | H3K36me3 | TRB1 | EMF1 | LHP1 | CLF/SWN |
| *icu11-5* | Up-regulated | Enrichment factor | 2.1 | 1.6 | 0.1 | 1.8 | 1.9 | 1.8 | 3.3 |
| | | Fisher's exact test (p-value) | 1.309e-35 | 9.433e-53 | 1.775e-49 | 2.022e-36 | 2.356e-23 | 2.020e-37 | 4.153e-27 |
| | Down-regulated | Enrichment factor | 3.5 | 1.8 | 0.1 | 0.7 | 2.8 | 3 | 3.6 |
| | | Fisher's exact test (p-value) | 1.848e-16 | 3.618e-10 | 2.986e-07 | 0.042 | 5.020e-10 | 7.706e-25 | 1.118e-04 |
| *icu11-5 cp2-1* | Up-regulated | Enrichment factor | 2.1 | 1.8 | 0.2 | 1.3 | 1.9 | 1.9 | 2.9 |
| | | Fisher's exact test (p-value) | 1.808e-160 | 0.000e+00 | 7.624e-184 | 2.979e-23 | 2.941e-106 | 1.201e-213 | 7.910e-94 |
| | Down-regulated | Enrichment factor | 1.6 | 1.6 | 0.8 | 0.9 | 1.6 | 1.5 | 1.5 |
| | | Fisher's exact test (p-value) | 4.350e-28 | 6.844e-106 | 9.150e-06 | 0.045 | 8.317e-28 | 3.405e-39 | 2.972e-06 |
| *emf2-3* | Up-regulated | Enrichment factor | 2.0 | 1.8 | 0.3 | 1.1 | 2 | 1.9 | 3 |
| | | Fisher's exact test (p-value) | 6.627e-96 | 5.331e-278 | 7.833e-98 | 2.964e-04 | 2.626e-101 | 6.249e-157 | 2.213e-75 |
| | Down-regulated | Enrichment factor | 1.7 | 1.7 | 0.5 | 1.1 | 1.8 | 1.5 | 1.8 |
| | | Fisher's exact test (p-value) | 6.914e-35 | 7.839e-167 | 4.799e-37 | 0.003 | 4.498e-46 | 1.521e-43 | 5.821e-12 |

Enrichment factors > 1 and < 1 indicate more and less overlap than expected of two independent gene lists, respectively. These data are complementary to those shown in Figure 4.

**Supplemental Table S4.** Primer sets used in this work

| Purpose | Oligonucleotide name(s) | Oligonucleotide sequences (5′ → 3′) | |
|---|---|---|---|
| | | Forward primer (F or L) | Reverse primer (R) |
| Genotyping | At1g22950_1F/R | ACCCTAACCTCTCAAACAAACCA | AGACTTTGTTAACCCAATCCGAC |
| | At1g22950_4F/R | CCTCTCAAACAAACCATCATCA | CGCTCAGTATCAGGGGAATATC |
| | SAIL_1215_B02_L/R | GAGCGATAACAGTGAGCTTGG | GACATTTTCAAACCATTCATGC |
| | AT5G51230_1F/R | TGTAATGGTTCAGAGATCAATAGAA | GTCCGTGCAATCTTGAGAATG |
| | LB1[a] | | GCCTTTTCAGAAATGGATAAATAGCCTTGCTTCC |
| RT-qPCR | qEARLI1_F/R | GATGCTCTCAGACTCGGTG | CGTCGAGGTCAACCAAACCT |
| | qRNS1_F/R | CGTTTTGGGAGCACGAATGG | ATCCCGGCTTTGGTTAGAGC |
| | qATPIRIN1_F/R | GAAGGAAGGTGAAGGAGCTG | TCTGTGAGGATGATCTGGGA |
| | qACTIN2_F/R | CACTTGCACCAAGCAGCATGAAGA | AATGGAACCACCGATCCAGACACT |
| | qAG_F/R | CCGATCCAAGAAGAATGAGCTCTT | CATTTTCAGCTATCTTTGCACGAA |
| | qSTK_F/R | TCAATCTCCCTTTTCTGCGCGTTT | TCAGGTCCAAGAAGCATGAGTTGC |
| | qSHP2_F/R | TCCGATCCAAGAAGCACGAGATGT | TCGTTTTGCAGCTCGATTTCCCTT |
| | qOTC_F/R | TGAAGGGACAAAGGTTGTGTATGTT | CGCAGACAAAGTGGAATGGA |
| Cloning[b] | TAP-CP2_Stop_F/R | ggggacaagtttgtacaaaaaagcaggctCTATGTCAAGTGAGCA GCGAGAAG | ggggaccactttgtacaagaaagctgggtTCAAGCTTGGGTTTGAC GTGG |
| | TAP-CP2_No Stop_F/R | ggggacaagtttgtacaaaaaagcaggctCTATGTCAAGTGAGCA GCGAGAAG | ggggaccactttgtacaagaaagctgggtCAGCTTGGGTTTGACGT GGTTTA |
| | TAP-ICU11_Stop_F/R | ggggacaagtttgtacaaaaaagcaggctTCATGTGCAATCAAAC TCCTCTTAG | ggggaccactttgtacaagaaagctgggtTTACTCGGCACATGATT TTGAAGC |
| | TAP-ICU11_No Stop_F/R | ggggacaagtttgtacaaaaaagcaggctTCATGTGCAATCAAAC TCCTCTTAG | ggggaccactttgtacaagaaagctgggtGCTCGGCACATGATTTT GAAGC |
| | TRB1_EYFP_Cter_F/R | gggacaagtttgtacaaaaaagcaggctTAATGGGTGCTCCTAAG CAGAAAT | ggggaccactttgtacaagaaagctgggtTCAGGCACGGATCATCA TTTTG |

**Supplemental Table S4 (continued)**. Primer sets used in this work

| Purpose | Oligonucleotide name(s) | Oligonucleotide sequences (5′ → 3′) | |
|---|---|---|---|
| | | Forward primer (F) | Reverse primer (R) |
| Cloning[b] | TRB3_EYFP_Cter_F/R | gggacaagtttgtacaaaaaagcaggctTAATGGGAGCTCCAAAGCTGAAG | ggggaccactttgtacaagaaagctgggtTTACCGAGTTTGGCTATGCATT |
| | LHP1_EYFP_Cter_F/R | gggacaagtttgtacaaaaaagcaggctTAATGAAAGGGGCAAGTGGTGCT | ggggaccactttgtacaagaaagctgggtTTAAGGCGTTCGATTGTACTTGA |
| | SWN_EYFP_Cter_F/R | gggacaagtttgtacaaaaaagcaggctTAATGGTGACGGACGATAGCAAC | ggggaccactttgtacaagaaagctgggtTCAATGAGATTGGTGCTTTCTG |
| | CLF_EYFP_Cter_F/R | gggacaagtttgtacaaaaaagcaggctTAATGGCGTCAGAAGCTTCGCC | ggggaccactttgtacaagaaagctgggtCTAAGCAAGCTTCTTGGGTCTA |

[a]Sequence taken from http://signal.salk.edu/tdnaprimers.2.html. [b] The *att*B1 and *att*B2 sequences are represented in lower case.

**Supplemental Table S5.** TAP and BiFC constructs

| Assay | Construct | Insert size (pb)[a] | Coordinates[b] | Destination vector |
|---|---|---|---|---|
| TAP | GSRhino-TAP-tagged ICU11 | 1,252 | Chr1:8125294-8127168 | pKCTAP |
| | GSRhino-TAP-tagged CP2 | 1,243 | Chr3: 6238266-6240396 | pKCTAP |
| BiFC | nEYFP-ICU11 | 1,255 | Chr1:8125291-8127168 | pSITE-nEYFP-C1 |
| | nEYFP-CP2 | 1,246 | Chr3: 6238263-6240396 | pSITE-nEYFP-C1 |
| | cEYFP-CLF | 2,769 | Chr2: 9955570-9960117 | pSITE-cEYFP-C1 |
| | cEYFP-SWN | 2,632 | Chr4: 886692-891473 | pSITE-cEYFP-C1 |
| | cEYFP-LHP1 | 1,399 | Chr5: 5827504-5829537 | pSITE-cEYFP-C1 |
| | cEYFP-TRB1 | 1,177 | Chr1: 1531806-1534305 | pSITE-cEYFP-C1 |
| | cEYFP-TRB2 | 1,132 | Chr2: 13732247-13734361 | pSITE-cEYFP-C1 |
| | cEYFP-TRB3 | 949 | Chr3: 18489451-18490731 | pSITE-cEYFP-C1 |

[a]The inserts were amplified from Col-0 cDNA using the *att*B primers shown in Supplemental Table S4. [b]TAIR10 Arabidopsis genome coordinates.

**Supplemental Table S6.** Quality control summary of RNA-seq analyses

| Sample[a] | Number of clean reads | Q30 bases (%)[b] | Read alignment rate (%) |
|---|---|---|---|
| Col-0 #1 | 53209306 | 95.78 | 98.81 |
| Col-0 #2 | 91762216 | 95.63 | 98.41 |
| Col-0 #3 | 61794508 | 95.58 | 98.51 |
| *cp2-1* #1 | 65083932 | 95.77 | 97.65 |
| *cp2-1* #2 | 73028616 | 94.71 | 98.26 |
| *cp2-1* #3 | 69311310 | 95.78 | 98.51 |
| *icu11-5* #1 | 65576476 | 95.77 | 98.39 |
| *icu11-5* #2 | 53564942 | 94.56 | 98.22 |
| *icu11-5* #3 | 61220290 | 95.52 | 98.09 |
| *cp2-1 icu11-5* #1 | 68890358 | 95.82 | 98.24 |
| *cp2-1 icu11-5* #2 | 75598942 | 94.34 | 98.16 |
| *cp2-1 icu11-5* #3 | 67551250 | 95.77 | 98.12 |
| *emf2-3* #1 | 55526214 | 95.45 | 98.38 |
| *emf2-3* #2 | 56487776 | 95.44 | 98.36 |
| *emf2-3* #3 | 66034456 | 94.22 | 98.39 |
| Col-0 inflorescence #1 | 56192392 | 95.61 | 98.16 |
| Col-0 inflorescence #2 | 55011404 | 95.78 | 98.29 |
| Col-0 inflorescence #3 | 54861298 | 95.56 | 98.30 |

[a]Each line corresponds to a different biological replicate. [b]The Q30 quality score indicates the percentage of bases whose correct base recognition rates are greater than 99.9% for the total bases.
